# mzQC: a versatile way to communicate quality information for biological mass spectrometry

**DOI:** 10.64898/2026.10.02.756144

**Authors:** Chris Bielow, Mathias Walzer, Nils Hoffmann, Julian Uszkoreit, Fatemeh Mirzadeh Sarcheshmeh, Franziska Nicolaus, Lars Andernach, Patrick Boschmann, Bo Burla, Eralp Dogu, Martin Eisenacher, Yasin El Abiead, David Jimenez-Morales, Steffen Neumann, Yasset Perez-Riverol, Timo Sachsenberg, Reza M. Salek, Michael Strobel, Stefan Tenzer, Tim Van Den Bossche, Olga Vitek, Mingxun Wang, David L. Tabb, Wout Bittremieux

**Author notes:** equal contribution (Chris Bielow, Mathias Walzer, Nils Hoffmann, Julian Uszkoreit).

## Abstract

Reliable quality control (QC) is essential for reproducible biological mass spectrometry, yet QC information is still typically stored in tool-specific, poorly interoperable formats that hinder comparison, reuse, and transparent reporting. We present mzQC, a community-developed standard from the HUPO Proteomics Standards Initiative for the compact, machine-readable representation of QC metrics and their associated metadata across mass spectrometry workflows. mzQC is implemented in JSON, organized to support both individual mass spectrometry acquisitions and collections thereof, with formal QC metrics definitions supported by controlled vocabularies to ensure unambiguous metric interpretation and semantic consistency. The format is supported by validation tools and reference libraries in Python, R, and Java, facilitating integration into existing software ecosystems. We demonstrate the utility of mzQC across diverse analytical modalities and QC settings, spanning longitudinal instrument monitoring, acquisition-method evaluation, study-level quality assessment, repository-scale data characterization, and spatially resolved mass spectrometry imaging. Together, these use cases show that mzQC provides not merely a file format, but a shared foundation for interoperable QC reporting, large-scale data assessment, and more rigorous, reproducible, and reusable mass spectrometry-based research.

## Introduction

Mass spectrometry (MS) has become an indispensable analytical technology in the life sciences, powering the detailed investigation of complex biological systems across domains such as proteomics, metabolomics, lipidomics, glycomics, and beyond. The potential of MS to evolve into a truly ubiquitous, robust, and high-throughput technology for clinical and translational research hinges critically on the reliability of the data it generates. Central to this reliability are the practices of quality assurance and quality control (QC), which ensure that data generated by MS experiments are accurate, reproducible, and interpretable.^1^ This is particularly important within the broader context of the reproducibility crisis in science,^2^ where unreliable experimental outcomes have highlighted the need for robust QC practices.

Quality assurance and quality control, in their essence, refer to the systematic processes and metrics used to assess the performance and reliability of an analytical system, focused on assessing experimental outcomes after data acquisition. In MS, this encompasses assessments of instrument performance, sample preparation consistency, and data acquisition fidelity.^1^ Without adequate QC, researchers risk drawing erroneous conclusions from their experiments. For example, variability in retention times or mass calibration errors might skew analyte profiles, while undetected sample carryover could compromise data interpretation. Instrument parameters— such as source pressure, ion injection times, scan frequencies, or temperature stability—can serve as valuable QC indicators, offering insights into operational consistency and early warning of potential deviations before they impact analytical results.^3^ Data quality issues not only jeopardize individual studies but also complicate meta-analyses and large-scale collaborative efforts, ultimately eroding the collective progress of the scientific community.

QC practices are broadly relevant across analytical disciplines, and their adoption in other fields demonstrates the significant impact of standardized practices. Genomics, for instance, has embraced standardized QC practices and reference datasets to ensure the reliability of sequencing results, fostering confidence and enabling significant advancements in the field.^4^ Similarly, MS has seen the emergence of standardized QC practices. In proteomics, various QC samples (low complexity “QC1” and high complexity “QC2” samples)^5^ are commonly used to monitor instrument stability and variability across experimental runs.^6^ Metabolomics often employs pooled samples or reference materials like the NIST SRM 1950 human serum,^7^ alongside blank samples to assess carryover and spiked-in compounds to evaluate detection limits.^8^ Community-driven initiatives, including those from the Association of Biomolecular Resource Facilities,^9^ the Metabolomics Quality Assurance & Quality Control Consortium,^10^ the Lipidomics Standards Initiative,^11^ and the Metaproteomics Initiative,^12^ have further advanced QC guidelines and benchmarks. These practices are indispensable for ensuring data quality, identifying sources of variation, and contextualizing experimental findings.

In parallel, an increasing number of bioinformatics tools have been developed to calculate QC metrics from MS data,^13^ offering researchers valuable insights into instrument performance and experimental consistency. These metrics make it possible to identify issues, to support decision-making during data processing, and to provide a basis for assessing experimental reproducibility. However, the utility of these tools is often hindered by the lack of standardization. Each tool employs its own custom file format, which, though suitable for internal workflows, leads to significant challenges when metrics need to be shared, aggregated, or interpreted externally. The lack of formal definitions and common representations limits interoperability, reduces transparency, and complicates efforts to compare or reproduce QC results. A standardized format with well-defined terms promotes rigor in metric generation, clarifies the meaning and interpretation of reported values, and enhances the utility of QC reports across software platforms and research groups.

To address this gap, the Human Proteome Organization Proteomics Standards Initiative (HUPO-PSI)^14^ Quality Control working group^15^ has developed mzQC, a file format specifically designed to unify the storage, communication, and verification of QC data in MS. The mzQC format provides a flexible, standardized, and machine-readable approach to QC, covering the diverse needs of biological MS across different omics domains and acquisition methods. Rather than prescribing judgments about data quality, mzQC offers an objective structure for reporting QC metrics alongside their contextual metadata. This supports transparent benchmarking, facilitates tool development, and enables consistent interpretation across studies. As MS continues to expand into new biological applications and analytical modes, mzQC establishes a critical foundation for advancing reliability, reproducibility, and community-driven innovation.

## Results

### Technical framework of mzQC

The mzQC format is specified in JavaScript Object Notation (JSON), a lightweight and widely supported data interchange format. This makes it easy to read and write mzQC data in most programming languages and provides a large amount of flexibility. The format employs a hierarchical schema to organize QC data, enabling storage of quality metrics and associated metadata both for individual mass spectrometry acquisitions (“`runQuality`”) and for collections of acquisitions (“`setQuality`”). This design ensures that the format is both expressive and machine-readable, allowing for easy integration into automated workflows and downstream analytical pipelines.

Central to mzQC’s design is the use of the PSI-MS controlled vocabulary,^16^ which provides standardized and unambiguous definitions for a wide array of QC metrics. These curated terms ensure consistent annotation and interpretation of QC data across software tools and experimental contexts. Additionally, the controlled vocabulary establishes semantic relationships among metrics to support structured querying, validation, and comparative analysis. While the PSI-MS controlled vocabulary provides the primary semantic framework for mzQC, the format also supports the use of additional controlled vocabularies, enabling domain-specific extensions and annotation of metrics beyond the currently defined PSI-MS terms. The PSI-MS controlled vocabulary is dynamically maintained, with new terms added on demand to accommodate emerging metrics and experimental applications.

The mzQC format also includes robust validation mechanisms. Syntactic validation is facilitated by its JSON Schema definition, ensuring that files conform to the expected structure. Semantic validation, which verifies the logical consistency and correctness of the data, is implemented in the pymzqc reference library.^17^ An online validator is available to easily check the syntactic and semantic validity of mzQC files without requiring specialized software installations.

To support adoption, user-friendly libraries for reading and writing mzQC files are available in Python, R, and Java, lowering the barrier to entry for researchers and developers.^17^ These libraries are designed to facilitate the creation, manipulation, and validation of mzQC files, streamlining the integration of QC practices into existing workflows. This broad software support ensures that mzQC is accessible to users with varying levels of computational expertise.

### mzQC for quality control of biological mass spectrometry

To illustrate the operation of mzQC for quality control in biological mass spectrometry, we present several use cases spanning proteomics, metabolomics, and lipidomics; targeted and untargeted workflows; data-dependent (DDA) and data-independent acquisition (DIA); and liquid chromatography coupled to tandem mass spectrometry (LC-MS/MS), gas chromatography coupled to mass spectrometry (GC-MS), and mass spectrometry imaging (MSI). Together, these examples demonstrate how mzQC can represent QC information across heterogeneous MS applications, from individual-run assessment and longitudinal instrument monitoring to spatially resolved and repository-scale analyses.

#### Identifying non-conforming mass spectrometry experiments using outlier detection

The first use case involves outlier detection in a proteogenomics study of *Mycobacterium tuberculosis*.^18^ This study investigated phenotypic variation across different *M. tuberculosis* strains, combining genomics and proteomics data to report novel strain-specific findings, including single amino acid variant peptides and a hypothetical protein derived from *de novo* assembled unmapped genomic reads. The proteomics component of the study consisted of 120 raw MS files acquired using DDA on a Thermo Scientific LTQ Orbitrap Velos instrument.

Confidence in these novel discoveries depends on high-quality MS data. We performed a QC analysis using different metrics and heterogeneous outlier detection methods to flag potentially problematic MS runs (**Figure 1a**). First, QuaMeter was used in its “ID-free” mode^19^ to calculate a set of 44 QC metrics directly from the raw MS data. Next, two multivariate outlier detection techniques were applied: robust principal component analysis (PCA)^19^ and the Local Outlier Probability (LoOP) algorithm.^20^ These techniques flagged a pair of MS runs that deviated significantly from the experiment’s overall data patterns. To complement the raw data-derived QC metrics, the PTXQC tool^21^ was used to calculate an additional set of metrics based on MaxQuant^22^ sequence database search results, incorporating peptide identification information. Using these metrics, LoOP was again employed to compute outlier scores. Together, these approaches provide a comprehensive and multifaceted perspective on data quality (**Figure 1b**). The outlier detection analysis revealed two MS runs that strongly deviate from the remainder of the dataset, likely due to potential electrospray ionization instability^23^ (**Figure 1c**). Notably, these two MS runs were measured at the end of an acquisition batch, with the instrument subsequently taken offline for maintenance (**Figure 1d**). Therefore, the deviations in instrument performance indicate that the results from these runs should be interpreted with caution, particularly in evaluating novel peptide and protein identifications. Additionally, a few other runs showed moderate outlier scores based on the MaxQuant-derived QC metrics (**Figure 1b**), suggesting that their identification results warrant careful examination.

**Figure 1.**
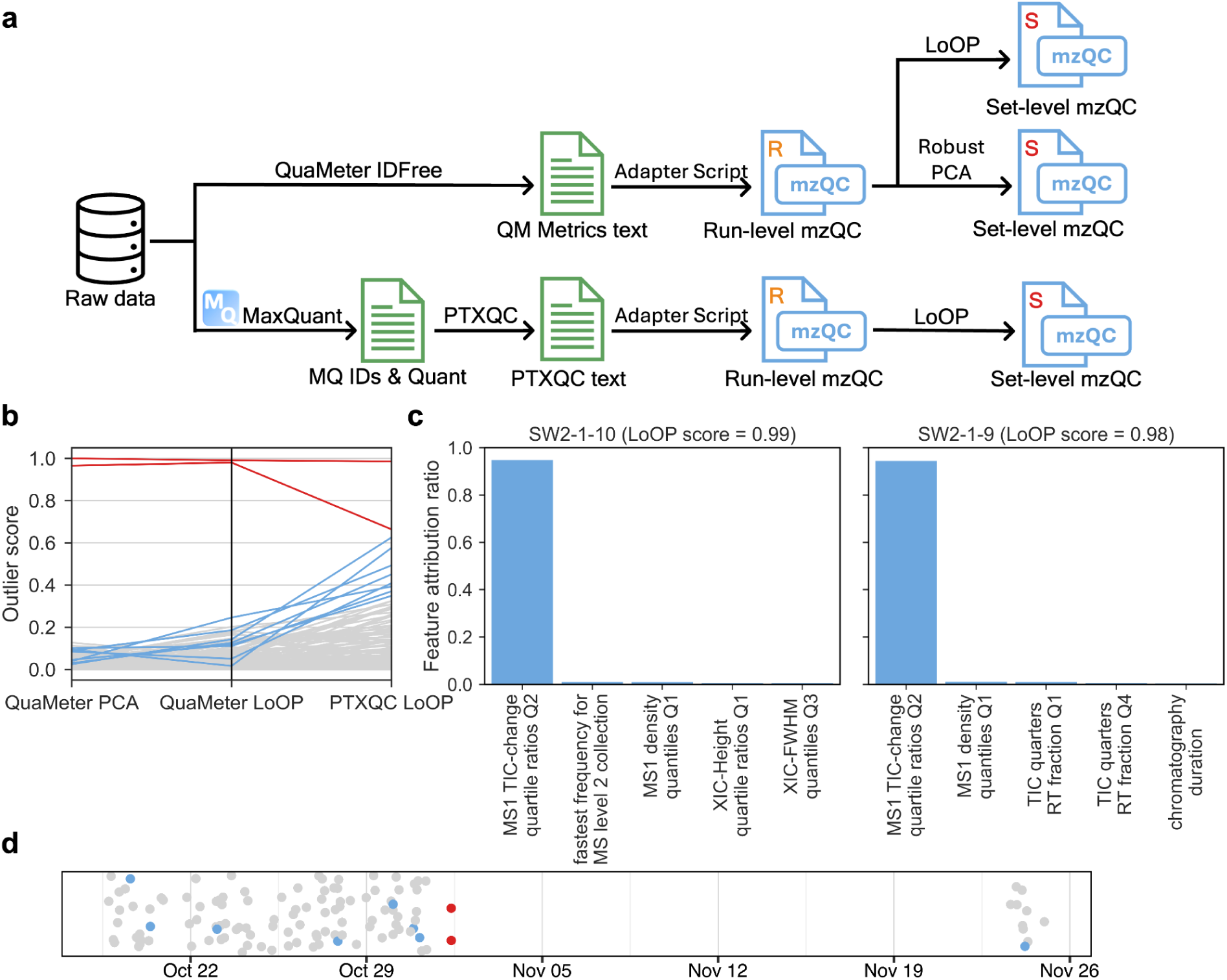
mzQC enables outlier detection to identify non-conforming MS runs. **(a)** QC workflow using mzQC as a unifying format. QC metrics were computed from raw MS data using QuaMeter and from MaxQuant search results using PTXQC. Run-level QC metrics were used as input to robust PCA and the LoOP outlier detection algorithm. Resulting outlier scores and QC metrics were aggregated into a single set-level mzQC file for downstream analysis. **(b)** Outlier scores for 120 MS runs show strong concordance across tools, with two runs flagged as outliers. QuaMeter- and PTXQC-derived metrics reveal complementary aspects of data quality. Runs exceeding an outlier score of 0.66 or 0.33 in any of the three analyses are colored in red and blue, respectively. **(c)** Detailed inspection of runs “SW2-1-9” and “SW2-1-10” indicates electrospray ionization instability as the primary anomaly, with substantial deviations in MS1 total ion current (TIC) contributing to high outlier scores. **(d)** Plotting runs by their start times reveals that the 120 experiments were measured in two batches; the two outlier experiments immediately precede the gap between batches.

The mzQC framework uniquely enables such a heterogeneous data quality evaluation. All calculated metrics for each MS run were stored in individual `runQuality` objects following analyses with QuaMeter and PTXQC. Next, the QC metrics served as input for the outlier detection algorithms, and the resulting outlier scores for the collection of MS runs were aggregated into a single mzQC file representing the set-level analysis. This workflow demonstrates mzQC’s ability to unify QC processes by providing a standardized format for both input and output across different tools and methodologies. The use of a controlled vocabulary ensures that QC metrics are machine-interpretable and unambiguously comparable, enabling straightforward integration of diverse computational approaches during data evaluation.

#### System suitability testing for longitudinal instrument performance monitoring

To demonstrate mzQC’s utility for system suitability testing, we analyzed longitudinal instrument performance data from the CPTAC Study 9.1, a coordinated multi-site effort by the Clinical Proteomic Tumor Analysis Consortium (CPTAC) to evaluate MS system reliability across four participating laboratories over approximately 6–8 months.^24^ During this period, each site executed repeated injections of standardized proteomic reference samples, amounting to several hundred QC runs per site, from which various metrics spanning retention time reproducibility, total peak area, and full width at half maximum (FWHM) were calculated to verify that instruments met predefined performance thresholds prior to experimental sample analysis. System suitability testing in this context serves as a critical mechanism for ensuring stable instrument operation and enabling reproducible data generation across time points and experimental batches.

We reprocessed the data from site 65 using Skyline^25^ and extracted and subsequently analyzed QC metrics using MSstatsQC,^26^ a software package developed for longitudinal QC in MS-based proteomics (**Figure 2**). MSstatsQC generates various statistical control charts, such as Shewhart and moving range control charts, to detect deviations from expected instrument behavior, revealing trends, drifts, or abrupt failures, and summarizing performance across analytes in interpretable statistical visualizations.

The mzQC format is a key enabler of this workflow, by providing a consistent, structured, and machine-readable way to store QC metrics across time. By capturing QC data in standardized mzQC files at each acquisition time point, longitudinal analyses become unambiguous, reproducible, and fully traceable. This consistency in data representation eliminates common sources of ambiguity such as changes in metric naming, data format drift, or undocumented transformations, all of which pose barriers to reliable long-term instrument monitoring. The need for such standardization is even more pronounced in multi-site studies like CPTAC, where harmonizing QC across diverse platforms, instrument models, and software ecosystems is essential. In this setting, mzQC’s design and use of controlled vocabularies ensures that QC metrics are both formally defined and semantically linked. This allows for unambiguous comparisons of performance across laboratories, as each site’s data are encoded using a shared, interpretable language. Without such formalism, comparisons risk being undermined by subtle inconsistencies in metric calculation, labeling, or interpretation.

**Figure 2.**
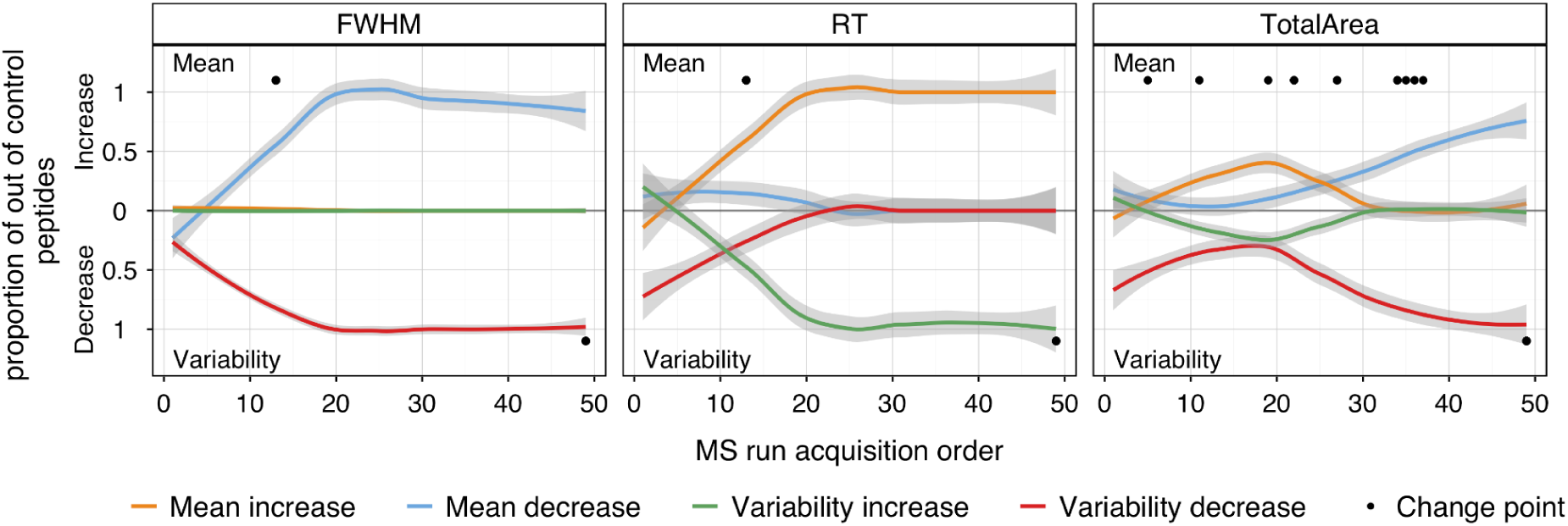
Longitudinal system performance monitoring using MSstatsQC control charts for CPTAC Study 9.1 (Site 65). Standardized proteomics QC samples were analyzed repeatedly over several months, peptide-level QC metrics were derived from Skyline results and stored in mzQC, and then used as input for downstream longitudinal analysis in MSstatsQC. The river plot shows an averages and range chart summarizing key QC metrics (e.g. FWHM, retention time, peak area) across multiple MS acquisitions. The vertical direction indicates the direction of change: upward trajectories represent increasing trends in the corresponding QC metric, whereas downward trajectories represent decreasing trends. Each curve represents the aggregated performance of a QC metric over time, averaged across all peptides. Colored lines indicate the proportion of peptides exhibiting deviations from expected values. Orange and blue lines represent positive and negative shifts in metric means, respectively, while green and red indicate increased or decreased variability, respectively. Black dots mark estimated time points where significant changes in performance occurred (change points). A gradual slope or increase in the number of change points indicates deteriorating instrument stability. This chart enables assessment of trends, drifts, and anomalies in instrument behavior over time.

#### Evaluating data-independent acquisition instrument methods

DIA has gained substantial traction in recent years due to its high reproducibility and identification comprehensiveness, making it a valuable strategy for large-scale proteomics. However, despite its growing adoption, DIA methods remain highly heterogeneous across laboratories.^27^ A central source of variation lies in the design of the isolation windows used to segment the precursor *m*/*z* range. For instance, some protocols use uniform window widths spanning the entire *m*/*z* range, while others employ variable-width strategies, allocating narrower windows to dense regions of the precursor space and wider windows to sparser regions. Some implementations, like staggered or overlapping windowing schemes, further increase method complexity. This diversity of configurations, often undocumented or inconsistently reported, creates substantial challenges for comparing or validating DIA experiments across sites and studies.

Such inconsistency undermines confidence in the comparability and reproducibility of DIA results, particularly in multicenter studies^28^ or method development efforts. Without standardized documentation and assessment of DIA parameters, it becomes difficult to evaluate whether observed differences in data quality are attributable to biological variation or to inconsistencies in method execution. Here, mzQC offers a solution by enabling transparent reporting and robust evaluation of DIA method performance.

To illustrate this, we applied DIAuditor, a QC tool developed to infer the DIA isolation window strategy used in a given experiment and to characterize the performance of the acquisition method via quality metrics. We analyzed 24 MS runs which employed a variable-window DIA method on a Thermo Q-Exactive mass spectrometer.^29^ This method used 19 isolation windows ranging in width from 24 to 222 *m*/*z*, with wider windows allocated to higher *m*/*z* regions to match the complexity of precursor ion distribution across the *m*/*z* space.

DIAuditor reports a range of metrics that characterize both the acquisition scheme and its effectiveness. One such metric is the total ion current (TIC) accumulated within each isolation window, computed by summing the TIC across all MS/MS scans assigned to that window (**Supplementary Figure 1**). DIAuditor captured a consistent relationship between isolation window mass range and TIC accumulation, with the lowest *m*/*z* window reaching 50% of its cumulative TIC by 34 minutes, on average, while the highest *m*/*z* window reached the same point at 92 minutes. This pattern, where low *m*/*z* windows accumulate intensity early in the run and high *m*/*z* windows accumulate intensity late in the run, reflects the longer retention and elution of larger peptides typical in reversed-phase liquid chromatography due to their greater potential for hydrophobic interactions. Deviations from this expected retention time–*m*/*z* relationship may indicate variability in chromatographic performance.

#### Repository-scale metabolomics QC reveals the analytical landscape of public mass spectrometry data

To demonstrate that mzQC is useful not only for study-level or instrument-level quality assessment, but also for large-scale characterization of public data resources, we performed a repository-scale QC analysis of 834,692 files^30^ from GNPS/MassIVE,^31^ MetaboLights,^32^ and Metabolomics Workbench^33^ by extracting a set of acquisition-related QC metrics for each MS file. mzQC provides a natural framework for this kind of heterogeneous information because it allows metrics to be represented in a uniform, machine-readable structure, irrespective of data origin.

These QC descriptors reveal the major classes of metabolomics experiments represented in public repositories. For example, metrics related to MS1 and MS2 ionization polarity separated files into broad analytical categories (**Figure 3a**), demonstrating that standardized QC metrics can recover the dominant structure of public metabolomics MS data directly from the measured properties of the files, without relying on often incomplete or inconsistent textual metadata. At repository scale, mzQC therefore does more than record quality information for individual files: it can provide an analytical index of the contents of an entire public data ecosystem.

**Figure 3.**
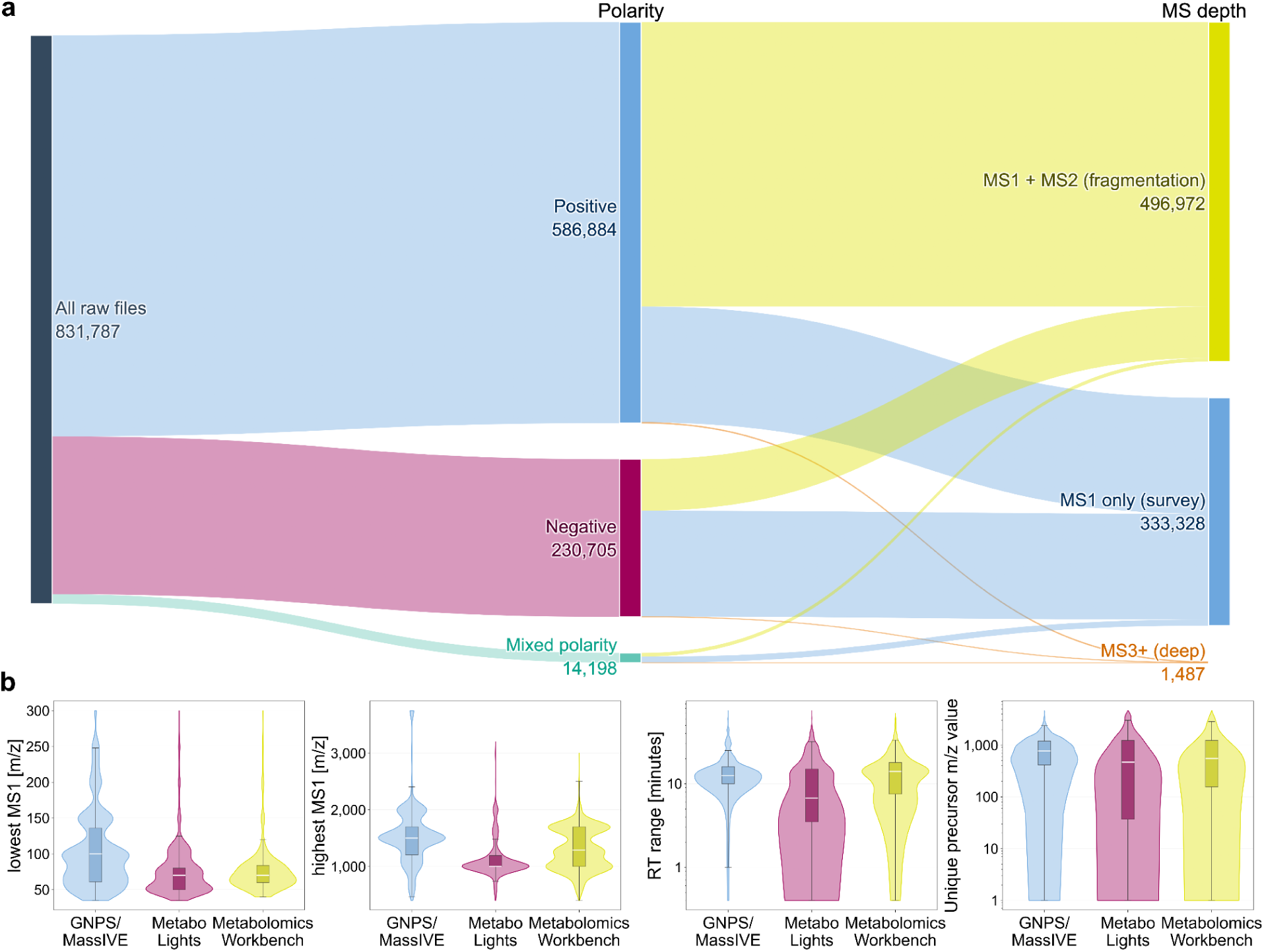
Repository-scale metabolomics QC landscape across public data resources encoded in mzQC. **(a)** Distribution of 831,787 public metabolomics files according to QC metrics capturing MS1 and MS2 ionization modes. These metrics separated the dominant acquisition classes observed in public metabolomics data, including positive- and negative-mode MS1-only experiments and positive- and negative-mode MS1/MS2 experiments. **(b)** Violin plots comparing the distributions of minimum MS1 *m*/*z*, maximum MS1 *m*/*z*, chromatographic retention time range, and the number of unique precursor *m*/*z* values per file across GNPS/MassIVE, MetaboLights, and Metabolomics Workbench, indicating repository-specific acquisition practices that may influence the observable chemical space and depth of MS/MS sampling.

The analysis also revealed repository-specific differences in acquisition design (**Figure 3b**), with potential consequences for the molecular classes captured and the suitability of datasets for secondary use. MetaboLights files generally had lower minimum and maximum MS1 acquisition boundaries than files from GNPS/MassIVE and Metabolomics Workbench. A lower minimum *m*/*z* may facilitate the measurement of small, polar metabolites, whereas a higher maximum *m*/*z* is better suited to larger secondary metabolites, lipids, conjugates, and other high molecular weight compounds. GNPS/MassIVE files also generally had longer chromatographic run times and contained more unique precursor *m*/*z* values. These characteristics are consistent with longer chromatographic separations and MS/MS-intensive acquisition strategies intended to maximize coverage of chemically complex samples.

Such differences are important for secondary analysis because repository selection can implicitly constrain the observable chemical space and the depth of precursor sampling. Public datasets from heterogeneous repositories should therefore not be treated as analytically interchangeable. Otherwise, benchmarking, meta-analysis, and machine learning studies may capture repository- or acquisition-specific signatures—such as mass-range boundaries, run duration, or precursor diversity—rather than the biological or chemical patterns of interest.

Repository-scale QC information also has immediate practical value for data discovery and reuse. Researchers seeking public datasets for benchmarking, reanalysis, or meta-analysis frequently need to identify experiments with specific analytical characteristics. When these characteristics are represented consistently in mzQC, they can be queried and compared directly across repositories, irrespective of local metadata conventions. This supports more informed dataset selection, more meaningful comparisons between experiments, and principled stratification of large public data collections according to acquisition design and basic performance characteristics.

The value of mzQC in this setting is therefore not limited to storage. At the scale of hundreds of thousands of files, QC information must be encoded consistently, interpreted unambiguously, and queried efficiently. mzQC is essential for this because it provides a shared representation for repository-scale QC metrics, links those metrics to defined semantic terms, and enables systematic filtering and aggregation across heterogeneous datasets. In this way, mzQC extends quality control from an internal laboratory practice into an infrastructure for the discovery, comparison, and reuse of public metabolomics data.

#### Quality assessment of targeted lipidomics data for reliable clinical measurements

MS-based lipidomics is increasingly used to characterize molecular changes associated with human health and disease, providing insights into disease mechanisms, biomarker candidates, and clinically relevant molecular signatures.^34^ The interpretation of lipidomics measurements in clinical studies depends on the ability to distinguish true biological variation from analytical variation introduced during sample preparation, instrument performance, and data processing. Comprehensive quality control is therefore essential to ensure that observed molecular differences reflect biological phenomena rather than technical artefacts and to support reproducible quantitative measurements across samples, batches, and studies.

To demonstrate the application of mzQC in this context, we captured quality information from a targeted lipidomics dataset with a comprehensive experimental design (**Supplementary Figure 2**)^35^ as standardized QC metrics generated using the MRMhub framework (**Supplementary Figure 3**).^36^ The exported metrics span complementary aspects of analytical performance, including quantitative reproducibility, separation from analytical background, and quantitative response behavior across the measured concentration range. Quantitative reproducibility was assessed using coefficients of variation calculated from pooled batch quality control samples, which monitor overall workflow performance, and pooled technical quality control samples, which primarily assess instrument performance and analytical stability. Technical quality control samples showed lower variability than batch quality control samples, with mean coefficients of variation of 14.9% and 22.6%, respectively (**Figure 4a**). Most monitored lipid features exhibited low variability, whereas a subset showed higher coefficients of variation, identifying features that may require additional evaluation during downstream analyses (**Supplementary Figure 4**).

**Figure 4.**
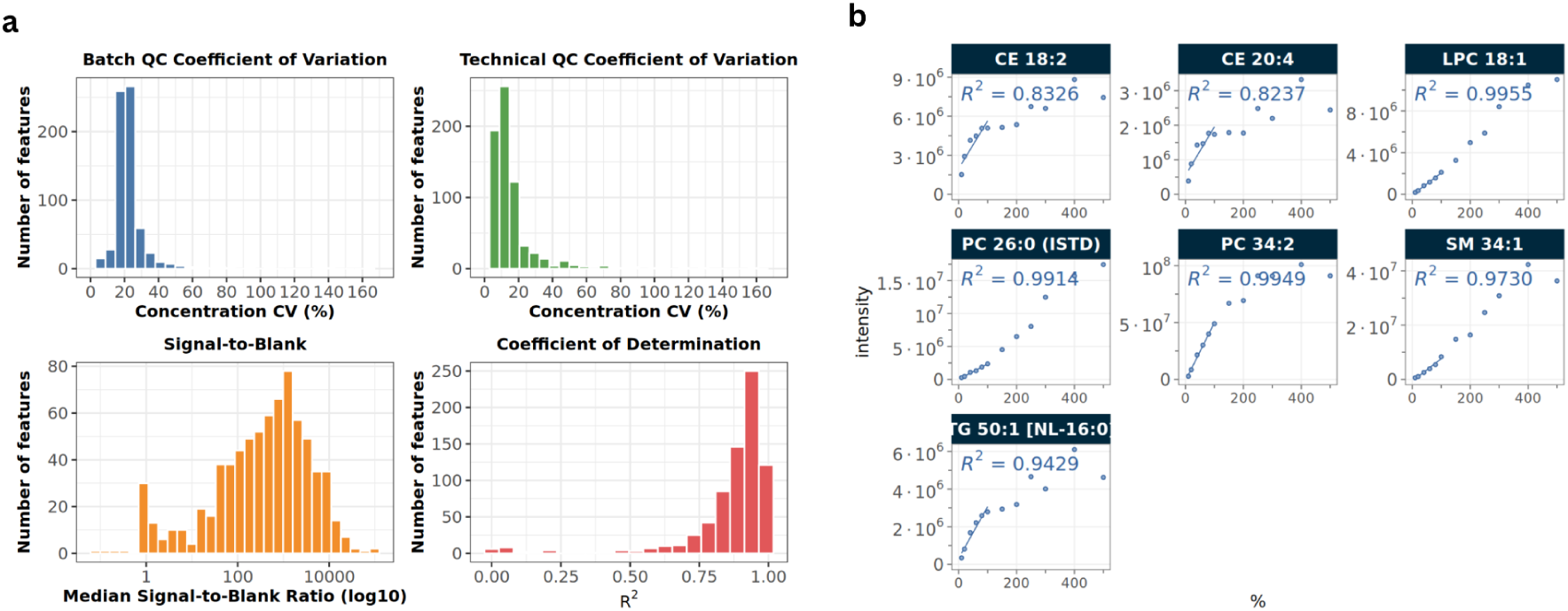
Standardized QC for targeted lipidomics using mzQC. **(a)** Distribution of exported QC metrics across monitored lipid species, including coefficients of variation for pooled batch quality control and pooled technical quality control samples, median signal-to-blank ratios, and response curve coefficients of determination (R²). **(b)** Examples of response curve measurements with different quantitative performance characteristics. Linear regression models fitted to response quality control dilution series measurements provided feature-specific coefficients of determination, slopes, and intercepts.

Signal-to-blank ratios were used to evaluate the separation of lipid feature signals from analytical background. Most lipid species showed signals substantially above procedural blank levels, while a subset exhibited lower signal-to-blank separation (**Figure 4a**). The broad distribution of signal-to-blank ratios reflects differences in analytical background contribution across lipid species. Quantitative performance was further assessed using response curve measurements obtained from dilution series experiments (**Figure 4b**). Most lipid features showed high response linearity, with coefficients of determination close to one, indicating consistent quantitative behavior across the analyzed concentration range. Features with reduced linearity represent candidates for further assessment of quantitative performance.

This evaluation of lipidomics data quality therefore relies on integrating multiple complementary quality control measurements rather than a single performance indicator. The workflow incorporated multiple types of QC samples, including pooled batch QC samples, pooled technical QC samples, procedural blanks, and response QC samples, each capturing different aspects of analytical performance. These measurements summarize groups of samples and QC metrics rather than individual MS acquisitions and were consequently represented as set-level quality information in mzQC. This enables heterogeneous quality control information generated throughout a clinical lipidomics workflow to be captured, exchanged, and reused in a standardized manner.

#### GC-MS quality control captures instrument degradation and maintenance effects

Standardized QC is also important for GC-MS, where changes in chromatographic performance and instrument response can accumulate over time. Dedicated QC mixtures containing compounds with known chromatographic and mass spectrometric behavior provide a practical means to monitor these changes, while representing the resulting measurements in mzQC extends the same standardized QC framework to GC-MS.

We evaluated longitudinal GC-MS performance during a measurement campaign targeting plant secondary metabolites using repeated measurements of a nine-compound reference mixture. Across 15 QC measurements acquired over approximately six months, reference compound intensities showed a progressive decrease during the first part of the campaign (**Figure 5a**). Instrument response increased following maintenance interventions, but subsequently declined again after several of the earlier interventions. The most pronounced recovery occurred after the final, more extensive maintenance procedure between QC measurements 14 and 15, which included shortening of the GC column.

The recovery in signal intensity after this intervention was accompanied by a systematic shift in chromatographic retention times. Median retention time deviation changed by approximately −6.1 s following column shortening (**Figure 5b**), consistent with the altered chromatographic path length. This illustrates that maintenance can improve one aspect of system performance while simultaneously changing another property relevant to downstream data interpretation. In mzQC-supported longitudinal studies, such shifts can identify batch boundaries and provide quantitative information for evaluating whether retention-time alignment or correction is required.

#### Spatially resolved QC metrics identify localized variation in MSI

MSI enables molecular characterization while preserving the spatial organization of biological samples, providing information about the localization of proteins, lipids, metabolites, and other molecular species within tissues.^37^ Unlike chromatography-based workflows, MSI datasets are organized according to spatial coordinates rather than acquisition time or chromatographic separation. Accordingly, spatially localized effects, such as heterogeneous matrix deposition, tissue-specific signal variation, or localized calibration deviations, can affect only part of an imaging experiment. mzQC’s data model was deliberately designed to flexibly represent QC information for various MS-based applications, including spatially resolved measurements generated by MSI.

**Figure 5.**
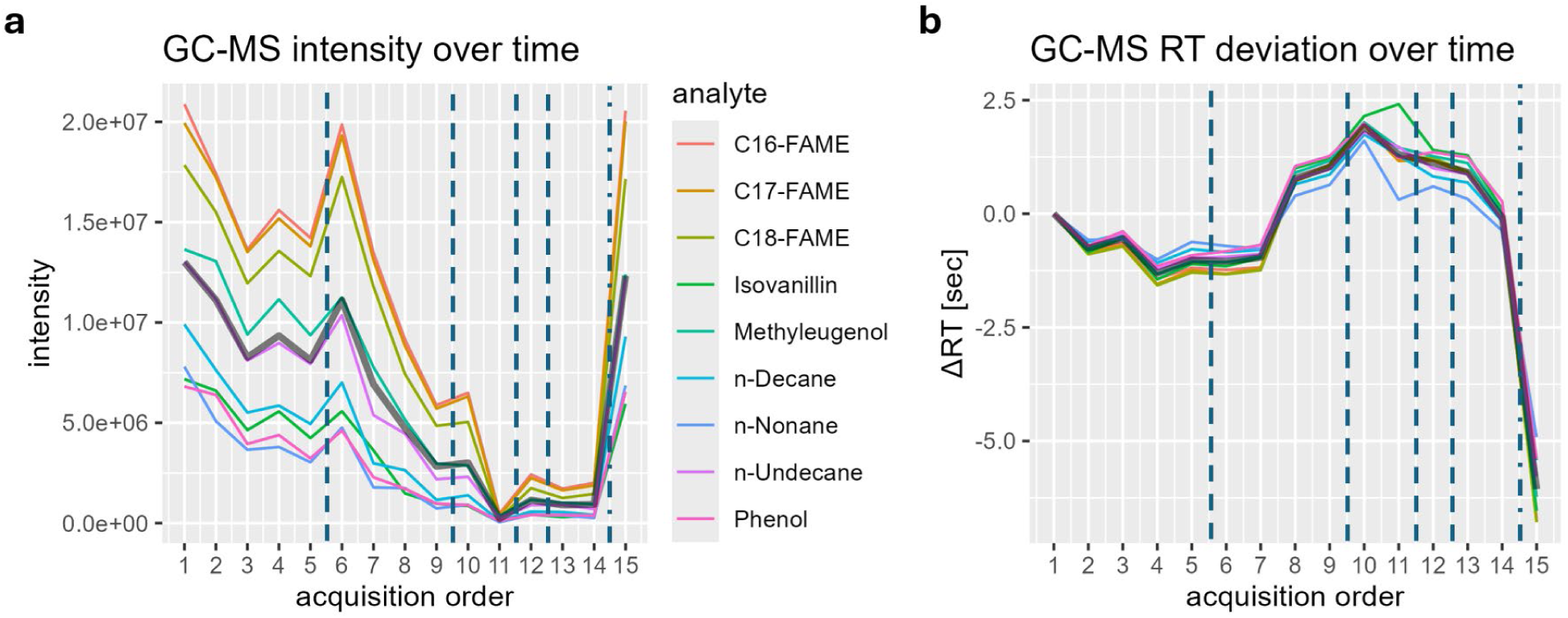
Longitudinal effects of instrument maintenance on GC-MS quality metrics. Maintenance between acquisitions is indicated as dashed line (minor maintenance) and dash-dotted line (major maintenance)**. (a)** Reference-compound intensities across sequential QC measurements, showing changes in instrument response over time and following maintenance interventions. **(b)** Retention time deviations relative to QC #1. Retention time deviations remained within the predefined acceptable range of ±2.5 s until maintenance before QC #15, which was associated with a systematic retention time shift of approximately −6.1 s.

We analyzed two publicly available MSI datasets: a matrix-assisted laser desorption/ionization (MALDI) lipidomics experiment of mouse brain tissue^38^ and a MALDI proteomics acquisition of primary cancer spheroids.^39^ In mzQC, spatially resolved MSI metrics can be represented as matrices in which each entry corresponds to a spatial position (pixel) in the imaging grid, while positions without acquired spectra can be represented as missing values. Both datasets demonstrated acquisition-specific spatial quality characteristics that could be captured using such matrix-valued QC metrics. Duplicate spatial coordinates were detected in both datasets, with 17 affected positions in the mouse brain dataset and 12 affected positions in the spheroid dataset. Inspection of the acquisition metadata revealed that these duplicates resulted from different acquisition behaviors: the spheroid dataset contained positions where two spatial columns were collapsed into a single coordinate, whereas the brain dataset contained additional spectra appended at the end of the acquisition and mapped back to an existing spatial position.

The spatial representation of QC metrics revealed quality patterns that were not apparent from global summary statistics alone. In the mouse brain dataset, the TIC displayed a strong spatial pattern caused by a polyethylene glycol (PEG)-related signal dominating the lower region of the image (**Figure 6a**). The corresponding base peak fraction, which measures the intensity fraction contributed by the most intense ion in each spectrum, showed spatial separation between tissue and matrix-dominated regions, with off-tissue areas exhibiting lower values due to the presence of multiple matrix-related ions (**Figure 6b**). Mass error likewise showed spatial variation, with a visible shift across tissue boundaries despite a relatively narrow global distribution (**Figure 6c**). The cancer spheroid dataset further demonstrated the importance of representing missing spatial positions explicitly. This experiment used a sparse acquisition strategy, with only a subset of positions in the imaging grid containing acquired spectra. The TIC and base peak fraction did not reveal substantial spatial artifacts in this dataset (**Figure 6d,e**). However, the mass error displayed a structured spatial pattern: the left region of the acquisition showed consistently higher mass errors than the right region (**Figure 6f**). This spatial shift was not apparent from the overall reference error alone, highlighting the importance of preserving the full spatial distribution of QC measurements.

**Figure 6.**
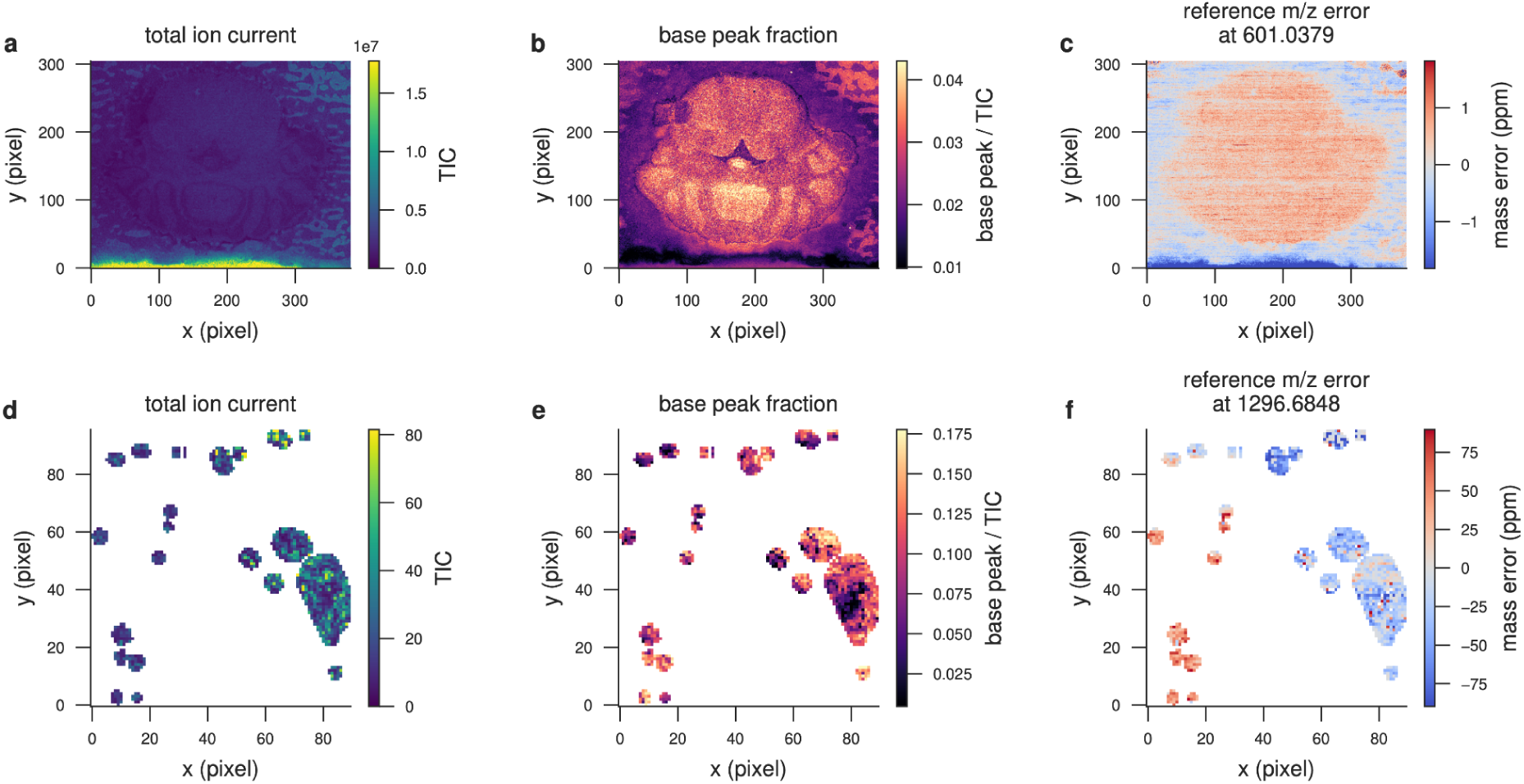
Spatially resolved QC metrics for MSI datasets represented in mzQC. (a–c) Mouse brain MALDI lipidomics dataset. **(a)** TIC image showing a strong PEG-related signal contribution localized to the lower region of the acquisition area. **(b)** Base peak fraction showing separation between tissue and matrix-dominated regions due to differences in dominant ion species. **(c)** Reference *m*/*z* error of the [4DHB−3H₂O+K]⁺ matrix cluster (601.0379 *m*/*z*), revealing a spatially structured mass shift across tissue boundaries. **(d–f)** Primary cancer spheroid MALDI proteomics dataset. White regions indicate positions without acquired spectra and are represented as missing values in the mzQC matrix metrics. **(d)** The TIC and **(e)** base peak fraction show no pronounced spatial quality deviations. **(f)** Reference *m*/*z* error of the Angiotensin I standard (1296.6848 m/z) reveals a spatially structured mass deviation, with consistently different mass errors between the left and right regions of the acquisition area.

Together, these examples demonstrate that MSI quality assessment requires QC representations capable of capturing spatially structured information rather than reducing measurements to single scalar descriptors. mzQC accommodates this requirement by supporting matrix-valued metrics alongside scalar and summary statistics. Combined with formally defined QC terms in the controlled vocabulary, this flexible representation enables mzQC to describe quality information across distinct mass spectrometry acquisition modalities, including spatially resolved imaging experiments.

## Conclusion

QC is a foundational component of any rigorous scientific discipline. In biological MS, which supports countless discoveries in proteomics, metabolomics, lipidomics, and systems biology, the need for robust, transparent, and interpretable QC is especially acute. MS experiments are highly sensitive to variations in sample preparation, instrument performance, and data processing. Without systematic QC, even minor deviations can propagate into misleading biological interpretations. While experimental QC practices are increasingly emphasized across MS-based omics, there has been a critical gap in standardized computational infrastructure to support these efforts.

To address this need, the HUPO-PSI Quality Control Working Group developed mzQC: a community-driven, open file format designed to unify the representation and exchange of QC data in biological MS. The format is implemented in JSON, offering flexibility and ease of adoption, while leveraging formally defined and semantically structured QC metrics maintained in a controlled vocabulary. We have demonstrated mzQC’s ability to support a broad range of QC-related tasks through representative use cases, including single-study quality assessment, longitudinal system suitability testing, repository-scale QC evaluation, and analysis of different acquisition strategies—such as untargeted DDA, DIA, targeted methods, and MSI—across multiple omics domains, including proteomics, metabolomics, and lipidomics.

Beyond improving reliability within individual studies or research groups, standardized QC information provides a foundation for large-scale data reuse and downstream analysis. It enables researchers and automated pipelines to evaluate whether datasets are fit for a particular analytical purpose, supports the harmonization of heterogeneous data collected across studies, and facilitates principled decisions about whether individual runs should be selected, weighted, or excluded. The boundary between metadata and QC information is often context-dependent: characteristics that describe an acquisition in one setting may serve as indicators of data quality in another. Acquisition parameters, instrument settings, or experimental design features can therefore provide essential context for interpreting QC measurements and assessing dataset suitability. mzQC supports this continuum by linking quality metrics with the contextual information required for their interpretation. These capabilities are particularly relevant to meta-analyses and machine learning applications, for which unrecognized technical variation or inconsistent dataset quality can introduce confounding signals and compromise model generalizability.

The utility of mzQC extends beyond its technical specification. It is supported by a growing ecosystem of software libraries for reading, writing, and validating mzQC files in Python, R, and Java.^17^ An online validator ensures syntactic and semantic correctness without requiring local setup. New QC metric terms can also be proposed through an open and documented process, allowing the standard and its controlled vocabulary to evolve in response to emerging technologies, acquisition strategies, and community requirements. Documentation, example datasets, and implementation resources available through the mzQC GitHub repository further support adoption and community participation.

Nevertheless, the practical value of a standard depends on the breadth and quality of its software support. QC-generating tools such as MsQuality,^40^ MSstatsQC,^26^ and OpenMS^41^ already support mzQC, and conversion scripts are available for outputs from tools including QuaMeter^42^ and PTXQC.^21^ However, broader native support will be required to establish a sufficiently rich ecosystem in which mzQC can be used consistently across instruments, analytical domains, and stages of data processing. The available Python, R, and Java libraries^17^ provide a basis for these integration efforts, but continued participation from software developers and instrument and workflow providers will be essential.

A further aspect is that mzQC does not prescribe a mandatory set of QC metrics or a single computational procedure for generating them. Instead, it provides a structure in which tools can store metric values and link them to controlled vocabulary definitions. Consequently, users and developers must ensure that each software implementation is consistent with the corresponding metric definition, including its calculation procedure, units, aggregation level, and required metadata. An mzQC file should therefore not be interpreted as intrinsically equivalent to every other mzQC file: its contents and analytical meaning depend on the metrics reported and on the software used to calculate them. Future work should include reference implementations, metric-specific validation datasets, and community benchmarking to verify consistency across tools.

Looking ahead, mzQC could become a core component of best practices in biological MS by supporting the standardized evaluation, comparison, and interpretation of MS data. As experiments become increasingly automated, large-scale, and distributed across laboratories, mzQC can provide the transparency and traceability needed to support robust scientific conclusions. Associating MS datasets with standardized QC files could strengthen laboratory practice by improving instrument monitoring, facilitating troubleshooting, and documenting data quality throughout an experiment. Because mzQC provides a consistent machine-readable representation of QC information, it also enables the development of user-facing interfaces such as graphical dashboards for interactive quality assessment and visualization. Combined with longitudinal QC data, these systems could support statistical trend modelling and early-warning approaches to detect gradual performance degradation before it affects experimental results. It could also support more systematic assessment of data quality during manuscript review, help readers evaluate published results, and enable researchers to identify public datasets that satisfy specific analytical requirements.

By enabling interoperable, extensible, and semantically defined QC reporting, mzQC provides infrastructure for more reliable and reusable MS data. Continued development of compatible software, reference implementations, and community-agreed metric definitions will determine the extent to which this potential is realized. Broad community participation will therefore be important for establishing QC as an integrated and transparent component of biological MS workflows.

## Methods

### mzQC data model and serialization

mzQC is serialized as JSON, using a hierarchical data model that separates file-level information, acquisition-level or set-level provenance, and individual quality metrics (**Figure 6**). The mzQC syntax is specified using JSON Schema and constrains the permitted elements, data types, cardinalities, and nesting relationships. An mzQC document contains a single top-level mzQC object; additional root-level properties are not permitted. The top-level object records the mzQC format version, an ISO 8601-compatible creation date and time (e.g., 2026-09-16T12:15:34Z), the controlled vocabularies used in the file, and at least one collection of either `runQuality` or `setQuality` objects. Optional file-level fields provide a description and contact information for the person or organization responsible for the file.

A `runQuality` object contains QC information derived from an individual MS acquisition, whereas a `setQuality` object contains QC information calculated for a defined collection of related acquisitions. Both object types implement the same `baseQuality` structure and contain two required components: metadata, which records the provenance and context of the QC analysis, and `qualityMetrics`, which contains the resulting QC measurements. Set-level metrics are intended to describe properties calculated across the defined collection of input files, rather than to provide an alternative container for unrelated run-level measurements. This distinction permits mzQC to represent both per-acquisition properties, such as the number of MS1 spectra or the chromatographic duration, and cohort-level results, such as an outlier score distribution, inter-run variability, or longitudinal performance summary.

The `metadata` object requires a unique and informative label, one or more `inputFiles`, and one or more `analysisSoftware` entries. Each inputFile records a unique file name, a URI identifying its location, and a controlled vocabulary term defining its file format. Optional file properties can describe additional attributes, such as checksums, native identifiers, or acquisition-related characteristics. Each `analysisSoftware` entry is represented by a controlled vocabulary accession and name together with the software version; a public URI for the software or its documentation can also be supplied. These fields provide provenance linking every QC result to the source data and software from which it was generated. Additional contextual information that is not part of the required metadata model can be included as controlled vocabulary parameters.

Individual QC results are stored as `qualityMetric` objects. A quality metric contains the accession and name of its corresponding vocabulary term, value, and unit. mzQC supports scalar values, ordered n-tuples, tables, and matrices. Tables are represented as named columns containing arrays of equal length, and the order and interpretation of columns and associated units follow the definition of the corresponding metric term.

**Figure 6.**
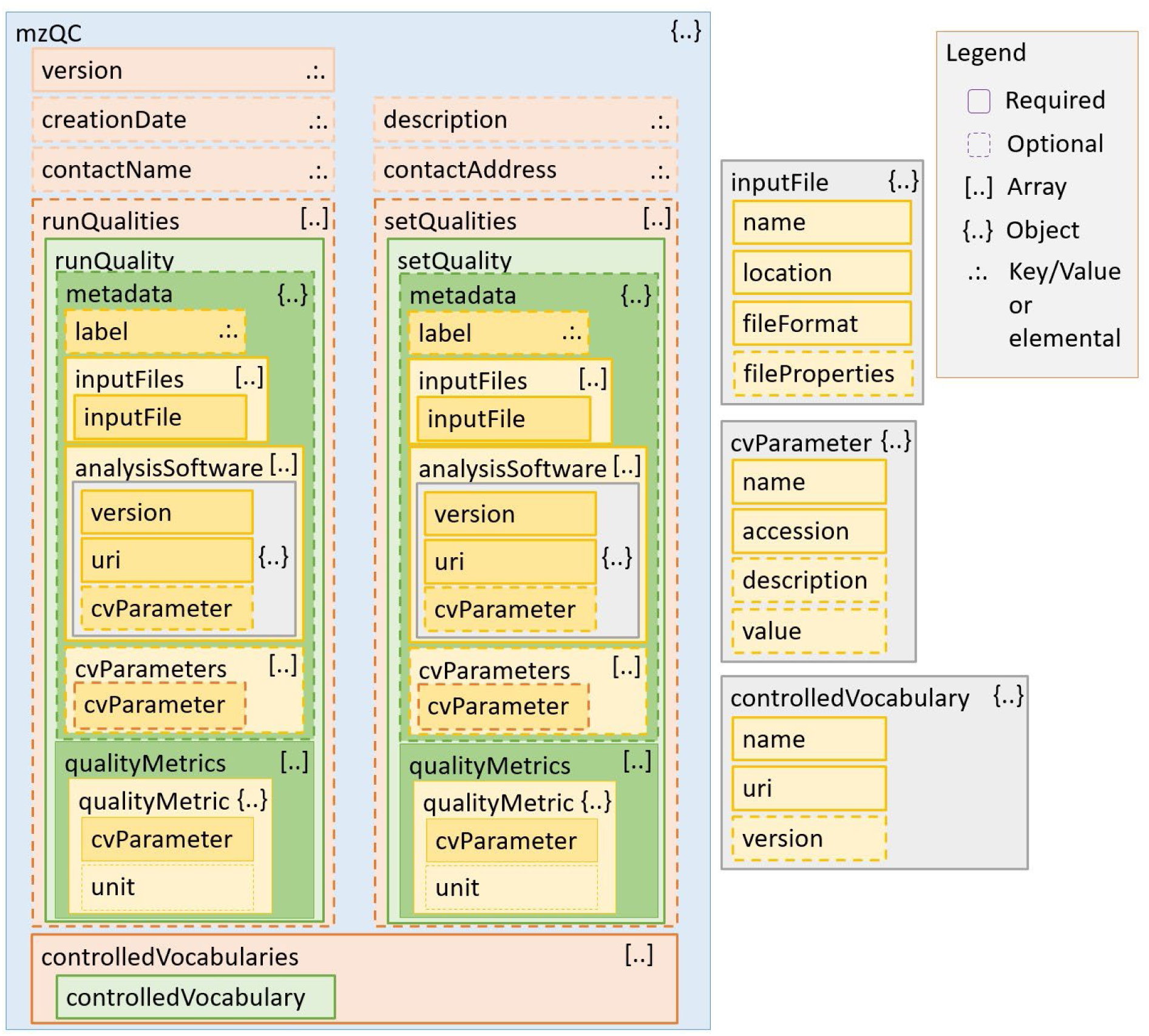
Schematic representation of the mzQC JSON schema. The mzQC format is organized as a hierarchical JSON structure with a top-level mzQC object containing file-level metadata, one or more `runQuality` and/or `setQuality` entries, and the controlled vocabularies used for annotation. Each `runQuality` or `setQuality` object stores contextual metadata, including input files, analysis software, and additional cvParameters, together with the reported `qualityMetrics`. Required and optional fields, as well as array and object relationships, are indicated to illustrate how mzQC supports structured representation of both run-specific and aggregate quality information.

The mzQC data model is intentionally descriptive rather than prescriptive. It defines how QC metrics, provenance, and derived QC analyses are communicated, but does not impose universal thresholds for acceptable instrument or data quality. Thresholds, reference ranges, statistical models, and pass/fail decisions can therefore be selected according to the analytical platform and experimental design, while the underlying measurements remain represented in a consistent and interpretable form.

### QC controlled vocabulary

The QC controlled vocabulary (QC CV) provides the semantic layer of mzQC. QC metrics are primarily sourced from the QC subset of the PSI-MS controlled vocabulary,^16^ indicated by the MS:4000XXX accession range. Each metric term has a stable accession, a preferred name, and a textual definition that specifies the quantity being measured and, where necessary, how it should be calculated or interpreted. The vocabulary also defines the expected value structure and unit or units of a metric. Consequently, two software tools that report the same accession are expected to produce semantically equivalent measurements, subject to any computational details stated in the term definition.

QC CV terms are classified through semantic relationships that describe relevant properties of the metric. These classifications can indicate, for example, whether a metric applies to a single acquisition or a set of acquisitions, and whether it describes MS1 data, MS2 data, chromatography, ion-source behavior, identification performance, or another aspect of the analytical workflow. Value types are represented explicitly as single-value, n-tuple, table, or matrix classes. Units are referenced from established ontologies where possible, including the Units of Measurement Ontology^43^ and statistical vocabularies.^44^ For structured metrics, the vocabulary defines the required columns, their order and interpretation, and any column-specific units. These relationships enable software to query metrics by analytical category and to validate their representation without relying on metric names or software-specific conventions.

The QC CV is extensible through a public term-request process via the HUPO-PSI Quality Control working group’s GitHub repository. Requests for new metrics require a name, an unambiguous definition, the intended mzQC representation and value type, and, where applicable, units, metric categories, and structured value columns. Proposed terms are reviewed by the mzQC working group before being incorporated into the controlled vocabulary. This process allows novel QC metrics to be supported without modifying the core mzQC JSON structure.

### Syntactic and semantic validation

mzQC validation is divided into syntactic and semantic components. Syntactic validation assesses conformance to the version 1.0.0 JSON Schema, including the presence of required elements, permitted property names, data types, cardinalities, URI and date-time formatting, and the organization of `runQuality`, `setQuality`, `metadata`, and `qualityMetric` objects.

Semantic validation assesses constraints that cannot be expressed completely in the JSON Schema. These checks included confirming that every declared controlled vocabulary is resolvable; that each accession and name pair occurs in one of the declared vocabularies; that a metric occurs only once within a given `runQuality` or `setQuality` object; and that each metric value and unit agrees with the value type and unit defined by its QC CV term. For table-valued metrics, all columns are required to have equal lengths. Labels are required to be unique within a file, and input-file locations are required to be unique within their parent quality object. Additional metric-specific plausibility checks, such as verifying the range or internal consistency of reported values, can be applied where defined.

The Python reference implementation, pymzqc,^17^ supports JSON serialization and deserialization, syntactic and semantic validation, file inspection, and merging of mzQC documents. The same validation functionality is exposed through the mzQC online validator (https://hupo-psi.github.io/mzQC/validator/), allowing files to be checked without a local software installation.

### Outlier detection in the *Mycobacterium tuberculosis* proteomics dataset

The analysis included 120 DDA MS runs from the *Mycobacterium tuberculosis* proteogenomics dataset, retrieved from the MassIVE database with accession MSV000081205 / PXD006843.^18^ Acquisition-level quality metrics were calculated from the Thermo RAW files using QuaMeter IDFree (version 1.1.21142).^19^ The resulting tab-delimited QuaMeter output was converted to an mzQC file using a custom Python script and the pymzqc library, mapping 44 QuaMeter output values to 17 scalar and n-tuple QC CV terms.

For the LoOP analysis, scalar QC metrics were represented as individual features and each element of an n-tuple metric was expanded into a separate feature. Features with variance below 0.01 were removed. Redundant features were then filtered sequentially by calculating absolute Pearson correlations and retaining only the first feature from pairs with a correlation greater than 0.9. The remaining features were robustly scaled by subtracting the median and dividing by the interquartile range. LoOP scores were calculated using 20 nearest neighbors, Euclidean distance, and a probabilistic local outlier factor normalization parameter of λ = 3.^20,45^ To identify the measurements contributing to an anomalous score, feature attribution values were calculated from the mean squared difference between each run and its 20 nearest neighbors. Contributions were normalized to sum to one across all retained features, and the five largest contributions were inspected for runs with LoOP scores greater than 0.66. QuaMeter-derived LoOP scores were represented as a set-level mzQC table linking each acquisition identifier to its outlier score.

A complementary robust PCA was performed in R using 16 QuaMeter-derived variables: the XIC50 fraction; the second components of the XIC FWHM, XIC-height, MS1 TIC-change, MS1 TIC, MS1-density, and MS2-density metrics; the sums of the second and third retention time fractions for TIC, MS1 scans, and MS2 scans; the numbers of MS1 and MS2 spectra; the fractions of MS2 precursors with charge states +2 and +3; and the maximum MS1 and MS2 acquisition frequencies. Pairwise Euclidean distances between runs were calculated using the first five principal components. The median distance from each run to all other runs was used as its robust PCA outlier statistic.^19^ These statistics were subsequently min–max scaled across the 120 runs to the interval [0,1] to facilitate comparison with the LoOP scores. The run identifiers and robust PCA scores were stored as a table in a set-level mzQC object.

For the identification-based analysis, MaxQuant (version 2.7.5)^22^ was run on 120 raw files using the default configuration and the *Mycobacterium tuberculosis* proteome from UniProt (UP000001584; SwissProt + TrEMBL; 3997 proteins). Identification-dependent QC metrics were obtained by applying PTXQC (version 1.1.6)^21^ to MaxQuant result tables. The PTXQC feature matrix was filtered using the same minimum variance threshold of 0.01 and absolute correlation threshold of 0.9 used for the QuaMeter analysis. Retained features were robustly scaled and LoOP scores were calculated with 20 nearest neighbors, Euclidean distance, and λ = 3. QuaMeter LoOP, QuaMeter robust PCA, and PTXQC LoOP results were joined by acquisition identifier. For visualization, each run was classified according to the largest of its three outlier scores: runs with a maximum score greater than 0.66 were classified as strong outliers, runs with a maximum score greater than 0.33 but not greater than 0.66 were classified as moderate outliers, and all remaining runs were left unflagged. Acquisition timestamps obtained from the QuaMeter output were used to order the runs chronologically and assess their positions relative to the interruption between acquisition batches.

### Longitudinal system suitability monitoring

Longitudinal system suitability data from Site 65 of CPTAC Study 9.1 were analyzed.^24^ The study comprised repeated measurements of a standardized proteomic reference sample acquired to monitor LC-MS/MS performance over time. Fifty system suitability acquisitions from Site 65 were analyzed, acquired between October and November 2011. They were measured as short blocks of consecutive injections interspersed with routine sample analysis. The analyzed acquisitions were a subset of the site’s system suitability series which was numbered up to 231 and no further runs were excluded before analysis.

Chromatographic peak detection and integration were performed in Skyline (version 24.1),^25^ monitoring doubly charged precursors for 15 reference peptides from 6 proteins. For each monitored precursor and acquisition, the extracted QC variables included the best retention time, chromatographic FWHM, peak start and end times, and total integrated peak area. Acquisition timestamps were obtained from the corresponding result files. Skyline-derived peak boundaries and quantitative measurements were used without further reintegration during subsequent mzQC conversion and longitudinal analysis.

The peptide-level results were exported to an mzQC file, where each acquisition was represented by a `runQuality` object containing input file provenance and acquisition time. Measurements for the monitored precursors were stored as a table-valued QC metric containing the precursor identifier, retention time, FWHM, peak boundaries, and total peak area.

Longitudinal QC analysis was performed using MSstatsQC (version 2.0).^26^ Upon import, measurements from all `runQuality` objects were combined into a single table, ordered by acquisition time, and mapped to the corresponding MSstatsQC variables. Individuals and moving-range control statistics were calculated separately for each monitored precursor and QC metric. The first five acquisitions were used as the guide set for estimating the reference mean and variability. Subsequent measurements were standardized relative to these guide values to identify positive or negative shifts in the metric mean and increases or decreases in temporal variability.

For each acquisition and metric, the river-plot analysis summarized the proportion of monitored precursors exhibiting each type of deviation. These proportions were smoothed across acquisition order to visualize gradual trends and abrupt changes in system performance. The plotted curves therefore represent the prevalence of peptide-level deviations rather than averages of the original retention time, FWHM, or peak area measurements. Change points were estimated independently for each precursor and metric after guide-set standardization. Changes in the mean were assigned to the acquisition position that best separated subsequent observations from the guide-set expectation, whereas changes in variability were detected using the corresponding moving-range model. The estimated change-point positions were overlaid on the river plot to indicate periods associated with systematic changes in instrument performance.

### Evaluating data-independent acquisition methods

Twenty-four DIA LC-MS/MS runs from a controlled-mixture proteomics dataset,^46^ available on MassIVE with accession MSV000081828, were analyzed. The data were acquired on a Thermo Scientific Q Exactive mass spectrometer using a variable-window method comprising 19 isolation windows across an MS/MS acquisition range of 400–1,220 *m*/*z*. The inferred window widths ranged from 24 to 222 *m*/*z*.

The mzML files were processed using DIAuditor (version 20250723 beta; https://github.com/dtabb73/DIAuditor). For each spectrum, the software extracted the MS level, scan start time, TIC, recorded peak count, mass resolving power, isolation window target *m*/*z*, lower and upper isolation offsets, and, when present, ion mobility information. DIAuditor generated both tab-delimited summary files and an mzQC document. The software does not require an externally supplied DIA window definition. Instead, MS/MS scans were grouped according to their isolation target and mobility value, and the lower and upper boundaries of each window were reconstructed by subtracting or adding the respective isolation offsets from the target *m*/*z*.

For each run and inferred isolation window, the window-specific TIC was calculated by summing the TIC reported for all assigned MS/MS scans. Scans were ordered by retention time, and cumulative TIC was calculated across the chromatographic run. The retention times at which the cumulative TIC first reached 25%, 50%, and 75% of the total window-specific TIC were recorded as TIC quartile retention times. The 50% value therefore represents the chromatographic time by which half of the signal acquired within that isolation window had accumulated. Additional window-level descriptors generated by DIAuditor included the number of MS/MS scans, retention time range, median acquisition cycle time, resolving power, and peak-count quartiles.

### Repository-scale metabolomics

QC metrics were generated for 834,692 mass spectrometry files obtained from three public metabolomics data resources: GNPS/MassIVE,^31^ MetaboLights,^32^ and Metabolomics Workbench.^33^ For each file, acquisition-level QC metrics were extracted using custom Python scripts as part of the Pan-ReDU project.^30^ These metrics were designed to summarize key analytical characteristics of each MS file, including acquisition mode, instrument sampling properties, and the observed chemical space.

The extracted metrics included the number of MS1 and MS2 spectra, stratified by ionization polarity; indicators of profile versus centroid acquisition; chromatographic retention time coverage; minimum and maximum MS1 *m*/*z* values; MS2 acquisition characteristics; and the number of unique precursor *m*/*z* values observed within each file. The minimum and maximum MS1 *m*/*z* values were used to characterize the measured mass range, retention time coverage was used as a proxy for chromatographic separation length, and the number of unique precursor *m*/*z* values was used to estimate the diversity and depth of MS/MS sampling. Together, these metrics provide an acquisition-level description of the analytical characteristics represented within each public repository.

To classify files according to major acquisition strategies, four scan-count metrics were selected for focused analysis: the number of MS1 and MS2 scans acquired in positive and negative ionization modes, respectively. These metrics were used to assign files to broad acquisition classes, including positive-mode MS1-only, negative-mode MS1-only, positive-mode MS1/MS2, and negative-mode MS1/MS2 experiments. Additional acquisition characteristics were compared across repositories, including minimum and maximum MS1 *m*/*z* range, chromatographic retention time range, and the number of unique precursor *m*/*z* values per file.

Metrics were extracted using custom Python scripts.^30^ For mzQC export, each file-level QC summary was encoded as a run-level mzQC object containing the calculated metrics and associated metadata.

### Clinical lipidomics quality assessment

QC data were generated using the MRMhub lipidomics workflow (version 0.9.9)^36^ developed by the Singapore Lipidomics Incubator (SLING). The dataset comprised 829 monitored features across 4,591 samples, including 460 identified lipid species, measured using targeted reversed-phase LC-MS/MS.^35^ Vendor-specific raw files were converted to mzML format using ProteoWizard msconvert (version 3.0.23277).^47^

Peak detection and integration were performed using the MRMhub-INTEGRATOR tool. Integration was performed using predefined feature and transition tables containing precursor– product ion transitions, expected retention times, and internal standard assignments, while feature-specific integration parameters were applied where required. The first two batch quality control samples were used as retention time reference anchors for chromatographic alignment.

Integrated peak areas were imported into MRMhub-QUANT together with sample annotations, feature information, interference correction information, internal standard assignments, and response curve metadata. The workflow performed sample outlier exclusion, interference correction, internal standard normalization, concentration quantification, drift correction, batch centering, and calculation of QC metrics. The resulting abundance profile consisted of detected lipid species and their absolute concentrations after these processing steps (**Supplementary Figure 5**).

QC metrics were calculated from pooled batch QC samples (N = 324), pooled technical QC samples (N = 325), procedural blanks (N = 5), and response QC samples (N = 36). Batch QC samples were generated from pooled aliquots of study samples and used to monitor overall workflow performance, including retention time stability, detector response, and measurement consistency. Technical QC samples were used to assess instrument performance and long-term analytical reproducibility. Procedural blanks were used to characterize analytical background contributions.

For each lipid feature, coefficients of variation were calculated separately for batch and technical QC samples as the standard deviation divided by the mean concentration across replicate measurements. Signal-to-blank ratios were calculated as the ratio between the median signal intensity in study samples and the median signal intensity in procedural blanks. Response curve parameters were calculated from linear regression models fitted to measured signal intensities across the response quality control dilution series for each lipid species. The coefficient of determination (R²), slope, and intercept of each regression model were extracted as QC metrics. QC metrics were exported as `setQuality`-based mzQC quality metric objects together with standardized metadata using the rmzqc R package.^17^

### Longitudinal quality control of GC-MS instrument performance

A QC mixture containing nine reference compounds was used: *n*-nonane, phenol, *n*-decane, *n*-undecane, methyleugenol, isovanillin, methyl hexadecanoate (C16-FAME), methyl heptadecanoate (C17-FAME), and methyl octadecanoate (C18-FAME). Compounds were selected to span the chromatographic gradient and to include different chemical functionalities, while the FAMEs additionally provided retention-index reference compounds. Concentrations were adjusted to 0.01 mg/mL.

Measurements were performed using a Shimadzu GC-2010 Plus gas chromatograph coupled to a Shimadzu GCMS-QP2010 Ultra mass spectrometer and equipped with a Restek Rxi-5Sil MS column (30 m × 0.25 mm internal diameter, 0.25 µm film thickness). Helium was used as carrier gas at 1 mL/min, with a purge flow of 3 mL/min. Samples (1 µL) were injected in splitless mode at 250 °C. The oven program was 50 °C for 1 min, followed by an increase from 50 to 330 °C at 25 K/min and a final hold at 330 °C for 2.8 min. MS settings were: interface temperature 250 °C, ion-source temperature 200 °C, electron ionization voltage 70 V, emission current 60 µA, scan speed 3333 Da/s, acquisition range m/z 50–450, threshold 150, and solvent cutoff 3 min.

Fifteen QC measurements were acquired over approximately six months, from 2025-09-25 to 2026-03-17. Maintenance was performed before QC measurements 6, 10, 12, and 13. A more extensive maintenance intervention was performed between QC measurements 14 and 15 and included shortening the analytical column by approximately 30 cm.

Vendor data were converted to mzML using ProteoWizard msconvert (version 3.0.25352)^47^ and processed using pyOpenMS (version 3.6.0.dev20260624).^48^ For each run, the workflow calculated the total ion chromatogram, applied Gaussian chromatogram smoothing, detected chromatographic peaks, and performed retention time alignment using dynamic programming, yielding a globally consistent, order-preserving one-to-one assignment.

Run-level QC metrics included the number of reference compounds detected and the median and interquartile range of retention-time deviation, signal-to-noise ratio, and chromatographic full width at half maximum. Metrics were exported for each acquisition as mzQC and visualized using rmzqc.^17^

### Mass spectrometry imaging QC metrics

QC metrics were generated for two publicly available MSI datasets. The first dataset consisted of a MALDI lipidomics experiment of mouse brain tissue (METASPACE https://metaspace2020.org/dataset/2025-04-27_13h26m30s).^38^ The sample was prepared using a 2,5-dihydroxybenzoic acid matrix deposited with a SMALDIPrep sprayer. Data were acquired using a Thermo Q Exactive Orbitrap mass spectrometer coupled to an AP-SMALDI5 AF ion source over an *m*/*z* range of 400–1600 using a 382 × 305 pixel grid with 25 μm spatial resolution. Data were exported in centroid mode as imzML files using RAW2IMZML (version 1.0r47).

The second dataset consisted of a sparse MALDI-TOF/TOF proteomics acquisition of primary cancer spheroids (PRIDE dataset identifier PXD013069) following trypsin digestion.^39^ The sample preparation, matrix deposition, acquisition parameters, and reference standards were as follows: α-cyano-4-hydroxycinnamic acid matrix and an internal standard mixture containing Angiotensin I, [Glu1]-Fibrinopeptide B, Dynorphin A, and ACTH clip 1–24 were applied using a Bruker ImagePrep station. Data were acquired on a Bruker ultrafleXtreme MALDI-TOF/TOF mass spectrometer over an *m*/*z* range of 800–4500 using a 90 × 96 pixel grid with 50 μm spatial resolution. Baseline removal and imzML export in profile mode were performed using SCiLS Lab (version 2016b).

Raw imaging data were imported using OpenMS (version 3.6.0.dev20260824).^49^ If multiple spectra were associated with the same spatial coordinate, only the first spectrum according to acquisition order was retained. Profile-mode spectra from the spheroid dataset were centroided using PeakPickerHiRes without signal-to-noise thresholding. All QC metrics were calculated from centroided spectra within the imaging geometry defined by the smallest bounding box containing all acquired coordinates.

Three spatial QC metrics were calculated independently for each acquired position. TIC was calculated as the sum of all peak intensities in a spectrum. Base peak fraction was calculated as the intensity of the highest-intensity peak divided by the TIC of the same spectrum. Reference mass error was calculated as the relative deviation between observed and theoretical *m*/*z* values, expressed in ppm. For reference mass error calculation, the highest-intensity peak within a predefined search window was selected. The [4DHB−3H₂O+K]⁺ matrix cluster (601.0379 *m*/*z*, ±20 ppm) was used for the mouse brain dataset, and the Angiotensin I reference peak (1296.6848 *m*/*z*, ±100 ppm) was used for the spheroid dataset. Reference mass definitions and search windows were stored as metadata associated with the QC metrics.

Each spatial metric was represented as a matrix corresponding to the imaging grid. Acquired positions were assigned metric values, whereas positions without spectra were explicitly represented as missing using null values. Null values were used instead of zero because zero may represent a valid measurement outcome for some metrics, such as signal intensity or perfect mass calibration. Each matrix metric was accompanied by an n-tuple summary statistic consisting of the first quartile, median, and third quartile calculated across non-null entries. Additional scalar QC metrics describing imaging geometry were calculated, including the x/y grid dimensions, acquired pixel fraction (number of unique acquired coordinates divided by total grid area), and duplicate pixel coordinate count (number of spatial coordinates associated with multiple spectra).

For each dataset, imaging-specific QC metrics were exported as mzQC files using pymzqc (version 1.0.1).^17^ Matrix-valued metrics were stored using the mzQC matrix representation, while associated summary statistics and acquisition descriptors were represented as scalar or tuple-valued metrics according to their definitions in the QC controlled vocabulary. For visualization, TIC images were clipped to the 99th percentile. Base peak fraction and reference mass error images were clipped to the first and 99th percentiles for display purposes. All spatial images were plotted using the lower-left coordinate as the origin to follow standard image coordinate conventions.

## Data availability

Mass spectrometry data analyzed in this study are publicly available:

- The *Mycobacterium tuberculosis* dataset used for outlier detection is available from MassIVE under accession MSV000081205 / PXD006843. MaxQuant result tables are available from MassIVE as reanalysis data (MassIVE RMSV000000714).
- Data for the longitudinal system-suitability analysis are available from Panorama at https://panoramaweb.org/cptac_study9.url.
- The DIA dataset is available from MassIVE under accession MSV000081828 and from PeptideAtlas under accession PASS00589.
- Repository-scale metabolomics data are available from GNPS/MassIVE, MetaboLights, and Metabolomics Workbench and were processed as part of the Pan-ReDU project.^30^
- Data for the clinical lipidomics analysis are available from Zenodo at DOI 10.5281/zenodo.21619042.
- GC-MS data are available from MetaboLights under accession MTBLS4827.
- The MSI data are available from the following:

- Mouse brain lipidomics imaging data are available from METASPACE under https://metaspace2020.org/dataset/2025-04-27_13h26m30s.
- Primary cancer spheroid imaging data are available from PRIDE under accession PXD013069. The “controlspheroids-nonormalization” acquisition was analyzed.

All derived mzQC files are available on Zenodo under the CC-BY-4.0 license at DOI 10.5281/zenodo.22959481.

## Code availability

The mzQC specification document, JSON Schema, example files, and user and developer documentation are available on GitHub under the CC-BY-4.0 license at https://github.com/HUPO-PSI/mzQC. Analysis scripts, derived mzQC files, and figure generation code related to the presented mzQC use cases are available as open source under the MIT license at https://github.com/MS-Quality-Hub/mzqc-use-cases. Source code for the clinical lipidomics analysis is available on GitHub under the MIT license at https://github.com/lifs-tools/mzqc-clinical-lipidomics-use-case.

Data processing, analysis, and visualization throughout this study were performed using a combination of computational tools, including Python packages such as NumPy,^50^ pandas,^51^ Matplotlib,^52^ seaborn,^53^ and Plotly,^54^ as well as additional software and libraries specified in the relevant Methods sections.

## Acknowledgments

M.Walzer acknowledges support from the H2020 EPIC-XS grant [Grant number 823839], BBSRC “Proteomics DIA” [BB/P024599/1], and from The Wellcome Trust [208391/Z/17/Z]. N.H. acknowledges support by ELIXIR, the research infrastructure for life science data, through activities in the ELIXIR Metabolomics Community and funding through 2024-SCIENCE-CMR. N.H. and P.B. acknowledge support from the de.NBI Cloud within the German Network for Bioinformatics Infrastructure (de.NBI) and ELIXIR-DE (Forschungszentrum Jülich and W-de.NBI-001, W-de.NBI-004, W-de.NBI-008, W-de.NBI-010, W-de.NBI-013, W-de.NBI-014, W-de.NBI-016, W-de.NBI-022). M.E., T.S., and D.L.T. acknowledge support by the Federal Ministry of Education and Research in the frame of de.NBI/ELIXIR-DE (M.E.: W-de.NBI-005, T.S.+D.L.T.: W-de.NBI-022), and S.N. W-de.NBI-011. Y.E. acknowledges support from the Austrian Academy of Sciences via APART-USA. T.S. acknowledges support by the Ministry of Science, Research and Arts Baden-Württemberg (LIBIS). S.T. acknowledges support from Bundesministerium für Bildung und Forschung (BMBF) as part of the National Research Node “Mass spectrometry in Systems Medicine” (MSCoreSys) [031L0217A/B] and the German Research Foundation (DFG, Deutsche Forschungsgemeinschaft; SFB 1292-3 Q1), as well as by the Research Center for Immunotherapy (FZI, Forschungszentrum für Immuntherapie) of the Johannes Gutenberg University Mainz. T.V.D.B. acknowledges support from the Research Foundation – Flanders (FWO) [1286824N]. O.V. acknowledges support from the Essential Open Source Software for Science award of the Chan Zuckerberg Initiative. M.Wang acknowledges the UC Riverside startup. D.L.T. was supported in part by a Strategic Health Innovation Partnership (SHIP) grant from the South African (SA) Department of Science and Technology (DST) and the SA Medical Research Council (SAMRC) to Gerhard Walzl. W.B. acknowledges support from the Research Foundation – Flanders (FWO) [G0AHY25N, W000924N].

Open Access Funding provided by Freie Universität Berlin.

We would like to thank the following contributors to the HUPO-PSI Quality Control Working Group meetings: Paul Brack, Jinmeng Jia, Axel Walter, and Weimin Zhu.

## Author contributions

**Mathias Walzer:** Conceptualization, Data curation, Investigation, Software, Supervision, Visualization, Writing – original draft, Writing – review & editing. **Chris Bielow:** Conceptualization, Data curation, Investigation, Methodology, Resources, Software, Validation, Visualization, Writing – original draft, Writing – review & editing. **Nils Hoffmann:** Conceptualization, Data curation, Investigation, Software, Supervision, Visualization, Writing – original draft, Writing – review & editing. **Julian Uszkoreit:** Conceptualization, Data curation, Software, Supervision, Writing – original draft, Writing – review & editing. **Fatemeh Mirzadeh Sarcheshmeh:** Data curation, Software, Validation, Visualization, Writing – review & editing. **Franziska Nicolaus:** Data curation, Software, Validation, Visualization, Writing – original draft. **Lars Andernach:** Investigation, Writing – review & editing. **Patrick Boschmann:** Data curation, Investigation, Software, Writing – original draft, Writing – review & editing. **Bo Burla:** Data curation, Investigation. **Eralp Dogu:** Data curation, Software, Writing – original draft, Writing – review & editing. **Martin Eisenacher:** Supervision, Writing – review & editing. **Yasin El Abiead:** Data curation, Investigation, Writing – review & editing. **David Jimenez-Morales:** Investigation. **Steffen Neumann:** Data curation, Writing – review & editing. **Yasset Perez-Riverol:** Software. **Timo Sachsenberg:** Investigation, Methodology, Software, Supervision, Writing – review & editing. **Reza Salek:** Conceptualization, Investigation. **Michael Strobel:** Data curation. **Stefan Tenzer:** Conceptualization, Writing – review & editing. **Tim Van Den Bossche:** Investigation, Writing – review & editing. **Olga Vitek:** Software, Writing – review & editing. **Mingxun Wang:** Data curation. **David L. Tabb:** Conceptualization, Investigation, Methodology, Project administration, Software, Supervision, Visualization, Writing – original draft, Writing – review & editing. **Wout Bittremieux:** Conceptualization, Funding acquisition, Investigation, Methodology, Project administration, Software, Supervision, Visualization, Writing – original draft, Writing – review & editing.

