## Supplemental Figures for "mzQC: a versatile way to communicate quality information for biological mass spectrometry"

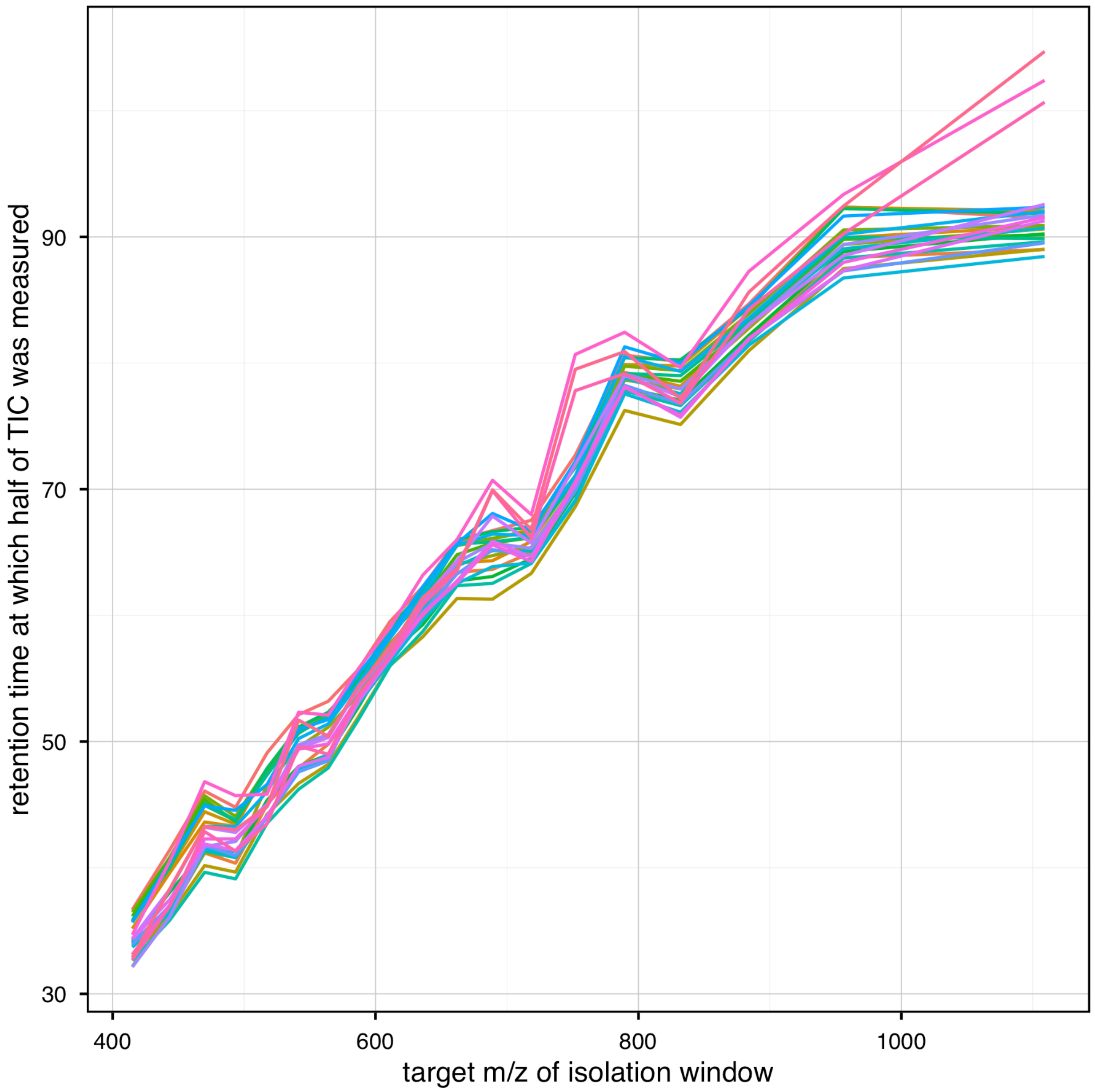


1. **Supplementary Figure 1. Relationship between isolation window *m/z* and retention time of ion accumulation in DIA experiments.** In DIA, higher *m/z* isolation windows tend to accumulate signal later in the chromatographic gradient. The plot shows the retention time at which each isolation window reaches 50% of its TIC, overlaid for 24 mzML files from dataset MSV000081828. Each line represents one DIA run. Lower *m/z* windows typically reach half of their TIC within the first third of the gradient, while the highest *m/z* windows require nearly the entire gradient to do so.

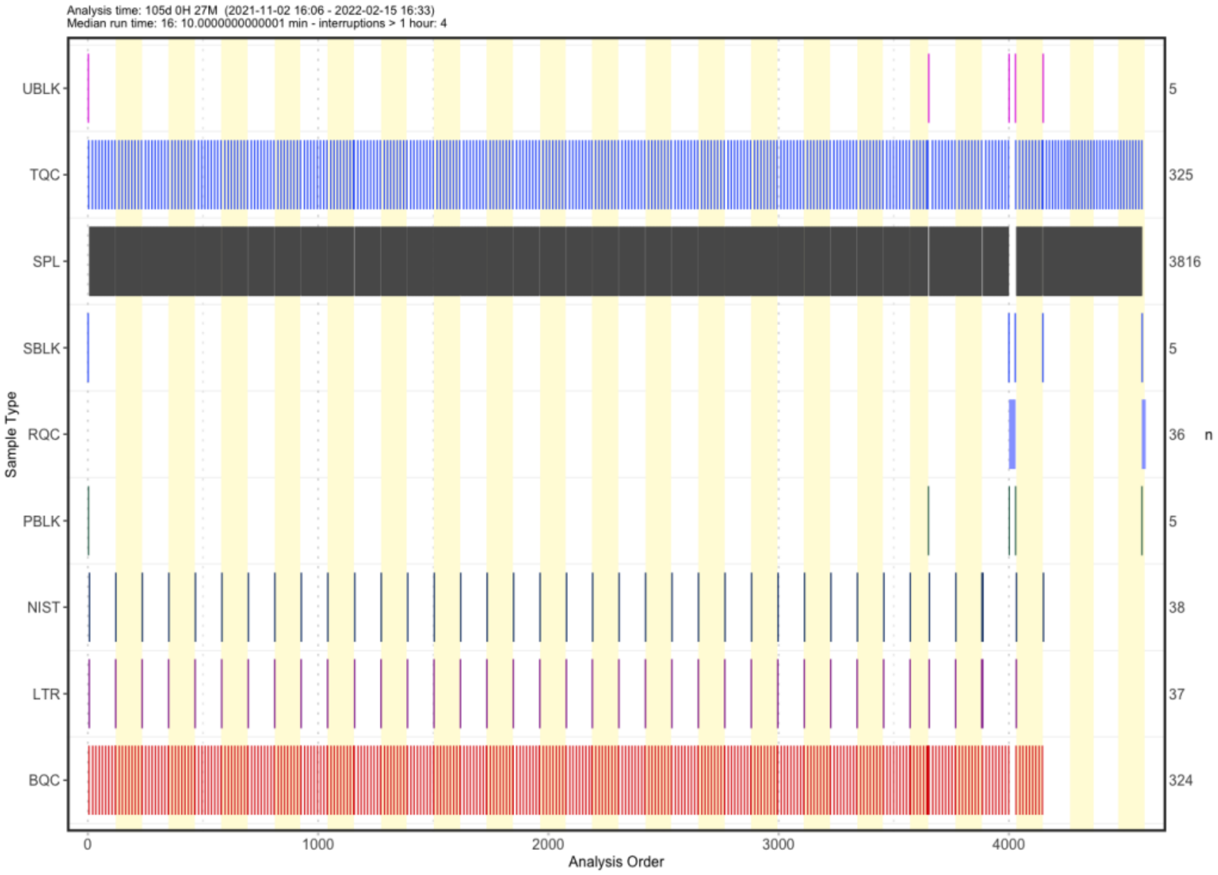


**Supplementary Figure 2. Experimental design and run sequence of the clinical lipidomics dataset.** Overview of the longitudinal targeted lipidomics experiment comprising more than 4,500 samples analyzed over 105 days. The figure shows the chronological arrangement of study samples, pooled batch QC samples, pooled technical QC samples, blanks, reference materials, and response QC samples throughout the analytical sequence. The inclusion and distribution of different QC materials enabled monitoring of analytical performance, background signals, instrument stability, and quantitative response across the complete measurement period. Abbreviations: UBLK, unextracted blank (no matrix, containing internal standard but not undergoing sample processing); TQC, technical quality control; SPL, study sample; SBLK, solvent blank/double blank (no matrix, no internal standard, containing solvents only); RQC, response quality control; PBLK, process blank (no matrix, processed with internal standard according to the sample workflow); NIST, NIST SRM 1950 standard reference material; LTR, laboratory-internal long-term reference standard; BQC, batch quality control.

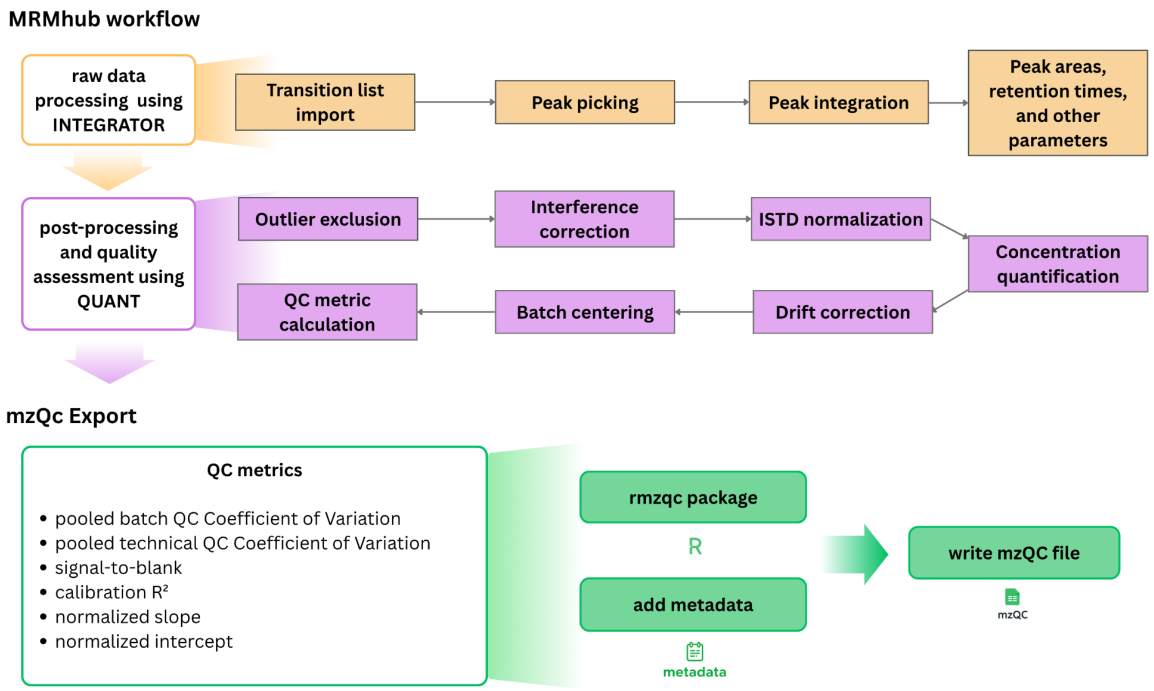


**Supplementary Figure 3. Overview of the MRMhub lipidomics workflow and mzQC export.** Peak integration results generated by MRMhub-INTEGRATOR were processed using MRMhub-QUANT, including outlier exclusion, interference correction, internal standard normalization, concentration quantification, drift correction, batch centering, and QC metric calculation. Selected QC metrics derived from multiple QC sample types were converted into mzQC quality metric objects and exported together with standardized metadata using the rmzqc R package.

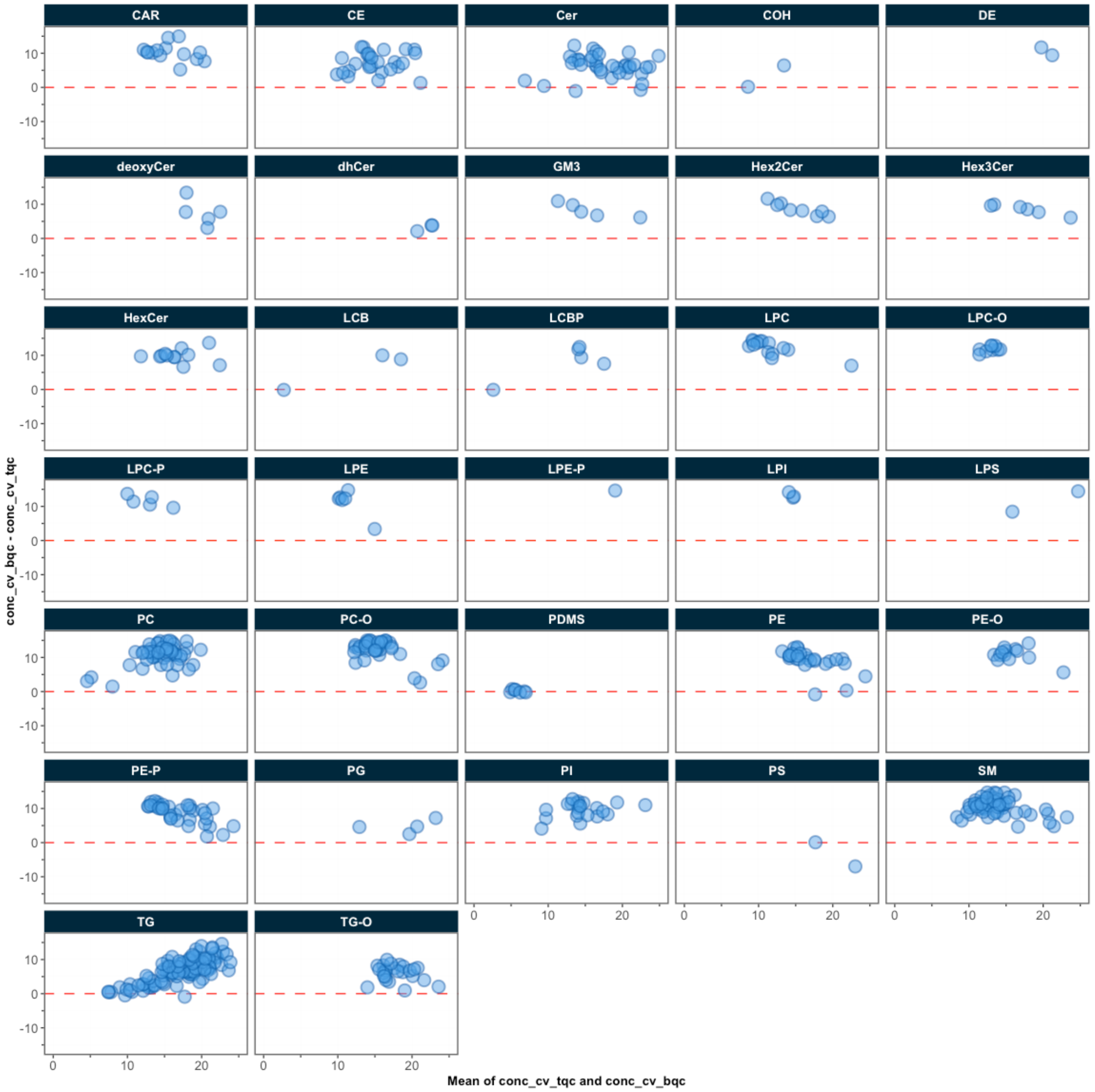


**Supplementary Figure 4. Comparison of concentration coefficients of variation between pooled batch QC and pooled technical QC samples for individual lipid features across lipid classes.** Each point represents a monitored lipid feature, while the panels correspond to lipid classes. The x-axis represents the mean coefficient of variation across batch and technical QC samples, and the y-axis represents the difference between batch and technical QC coefficients of variation. Coefficients of variation from pooled batch QC samples capture variability across the overall analytical workflow, including sample preparation and measurement steps, whereas coefficients of variation from pooled technical QC samples primarily reflect analytical measurement variability. Thus, positive values on the y-axis indicate a higher coefficient of variation in batch QC samples, whereas negative values indicate a higher coefficient of variation in technical QC samples. The majority of lipid features exhibit higher coefficients of variation in pooled batch QC samples than in pooled technical QC samples, indicating that additional sources of variability are captured when assessing the complete workflow performance.

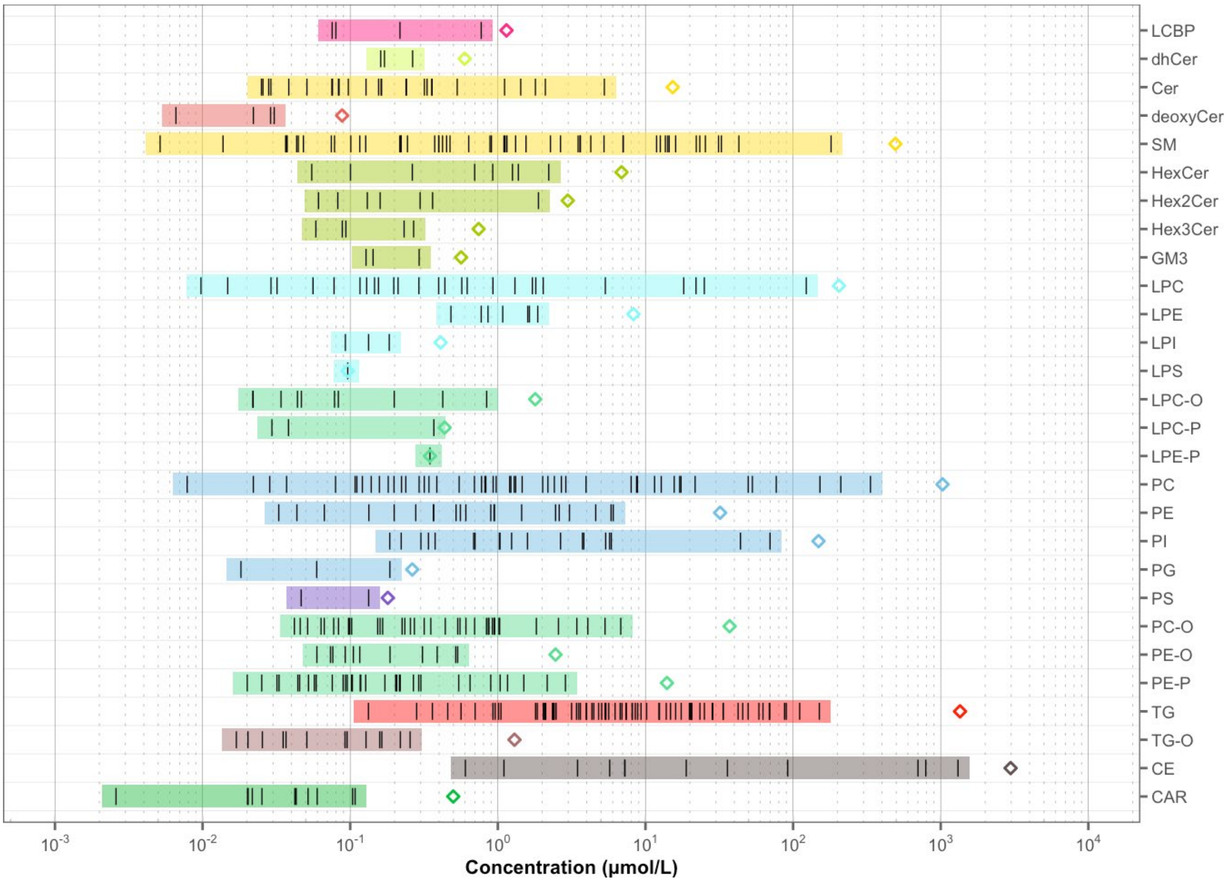


**Supplementary Figure 5. Quantitative abundance profile of detected lipid species after data processing.** Distribution of absolute lipid concentrations (μmol/L) across detected lipid species after feature filtering, normalization, quantification, drift correction, and batch correction. The abundance profile spans approximately six orders of magnitude, reflecting the broad dynamic range of lipid concentrations measured in the dataset.
